# Identification of *FoxP* circuits involved in locomotion and object fixation in *Drosophila*

**DOI:** 10.1101/2020.07.15.204677

**Authors:** Ottavia Palazzo, Mathias Raß, Björn Brembs

## Abstract

The *FoxP* family of transcription factors is necessary for operant self-learning, an evolutionary conserved form of motor learning. The expression pattern, molecular function and mechanisms of action of the *Drosophila FoxP* orthologue remain to be elucidated. By editing the genomic locus of *FoxP* with CRISPR/Cas9, we find that the three different *FoxP* isoforms are expressed in neurons, but not in glia and that not all neurons express all isoforms. Furthermore, we detect *FoxP* expression in, e.g., the protocerebral bridge, the fan shaped body and in motorneurons, but not in the mushroom bodies. Finally, we discover that *FoxP* expression during development, but not adulthood, is required for normal locomotion and landmark fixation in walking flies. While *FoxP* expression in the protocerebral bridge and motorneurons is involved in locomotion and landmark fixation, the *FoxP* gene can be excised from dorsal cluster neurons and mushroom-body Kenyon cells without affecting these behaviors.

## 1. Introduction

The family of *Forkhead Box (Fox*) genes comprises a large number of transcription factors that share the evolutionary conserved forkhead/winged-helix DNA-binding domain [1]. In mammals, the *FoxP* subfamily (*FoxP1-4*) members [2] are abundantly expressed during the development of multiple cell types, such as cardiomyocytes, neurons, lung epithelial secretory cells and T-cells [3]. In particular, *FoxP1* and *FoxP2* have generated interest because of their roles in regulating the development of cognitive processes such as speech and language acquisition [4–13].

Humans with *FOXP1* deletions present with mild mental retardation, delayed onset of walking, gross motor impairments and significant language and speech deficits [8]. Mutations in *FOXP2* cause a severe speech and language disorder characterized by deficits in language processing, verbal dyspraxia and impaired grammatical skills, without affecting other traits severely [4,5]. The function of *FoxP* genes in vocal learning appears to be evolutionary conserved as knockouts of the zebra finch orthologue of human *FOXP2* during the critical song learning period alters the structure of the crystallized song in the adults [14]. Such vocal learning is a form of motor learning that proceeds slowly from highly variable ‘babbling’ (in humans) and ‘subsong’ (in zebra finches) towards more stereotypic language and crystallized song, respectively. This specific kind of learning has been classified as a form of operant learning [15–17]. It was recently shown that, as in humans and zebra finches, also in flies, *FoxP* is involved in such operant learning [18].

The original *forkhead (fkh*) gene was identified in the fruit fly *Drosophila melanogaster [19],* where mutations cause defects in head fold involution during embryogenesis, causing the characteristic “fork head”. In contrast to chordates with four *FoxP* family members, only one orthologue of the *FoxP* subfamily is present in flies (*dFoxP).* The *dFoxP* gene gives rise to three different transcripts by alternative splicing [2,18,20]: *FoxP-isoform A (FoxP-iA), FoxP-isoform B (FoxP-iB*) and *FoxP-isoform IR* (Intron Retention; *FoxP-iIR*) (Fig. 1A). The currently available reports as to the expression pattern of the *FoxP* gene have been contradictory and nothing is known as to whether the different isoforms are differentially expressed in different cell types. To resolve these issues we have tagged the endogenous *FoxP* gene, analyzed the isoform-specific expression patterns and compared them with the expression of a selection of cell-type specific markers.

**Fig. 1:**
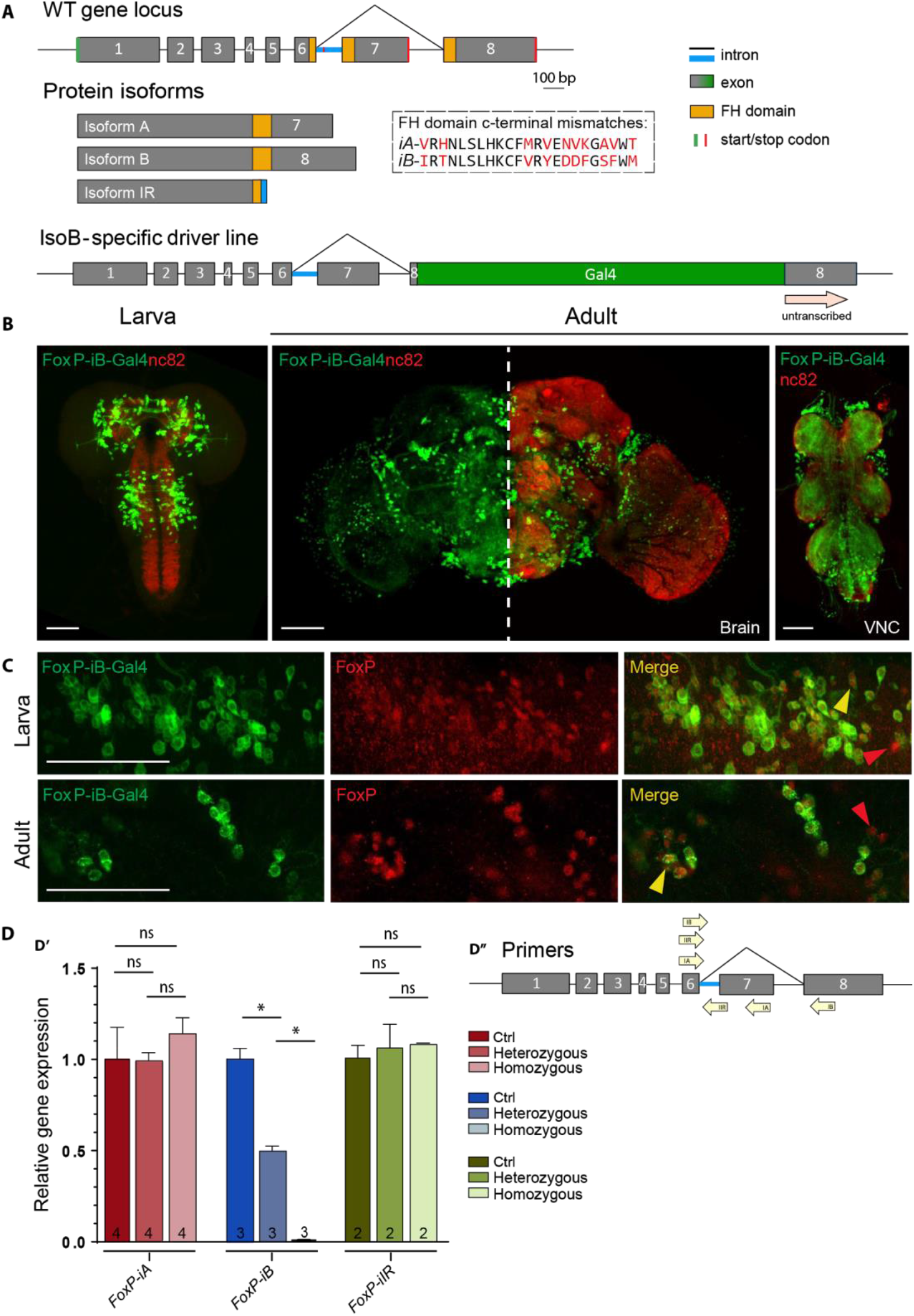
*FoxP-iB expression in the* Drosophila *nervous system.* (A) Schematic representation of the *FoxP* gene locus before (above) and after (below) insertion of a Gal4 sequence into exon 8. (B) *FoxP-iB-Gal4>CD8-GFP* expression pattern costained with nc82 in 3rd instar larvae, adult brain and adult VNC. (C) Driver line costained with a polyclonal FoxP antibody in larval and adult brain. The yellow arrowheads indicate colocalization, while the red ones indicate cells only positive for the antibody staining. (D’) RT-qPCR for FoxP-iA, iB and IR on controls and hetero and homozygous *FoxP-iB-Gal4* mutant. (D’’) Primers used for the RT-qPCR. Data are expressed as means ± SEM. *p<0.005. Scale bars: 50 μm.

Flies with a mutated *FoxP* gene not only show impairments in operant learning but also in motor coordination and performance of inborn behaviors [18,20–22]. While isoform-specific alleles did show different phenotypes as well as different degrees of severity of these impairments, it remains unknown which neurons require *FoxP* expression at what developmental stage(s) for normal locomotor behavior. We therefore knocked out the *FoxP* gene in a spatiotemporally controlled manner and analyzed spatial and temporal parameters of locomotor behavior in the resulting mutants in Buridan’s paradigm.

## 2. Materials and Methods

### 2.1. Fly strains

Fly stocks were maintained at 18 °C (Table 1). Before experimental use, flies were kept at 25 °C, in a 12/12 hours light/dark regime at 60% relative humidity for at least one generation. All crosses were raised at 25 °C (except for the ones involving the temperature sensitive Gal4 inhibitor Gal80^ts^ *[23,24]* that were raised at 18 °C) using 4-6 females and 2-4 males. For expression pattern visualizations, the *FoxP-iB-Gal4* and *FoxP-LexA* driver line, respectively, were crossed with the appropriate effector lines containing different GFP or RFP variants (Table 1). For behavioral analysis involving the *UAS-t:gRNA(4×FoxP*), this effector line was always first crossed with a *UAS-Cas9* line, and the resulting double-effector offspring with the appropriate driver line for each experiment (*ELAV-Gal4, D42-Gal4, C380-Gal4, cmpy-Gal4, ato-Gal4* and *ELAV-Gal4;Tub-Gal80^ts^*).

**Table 1:**
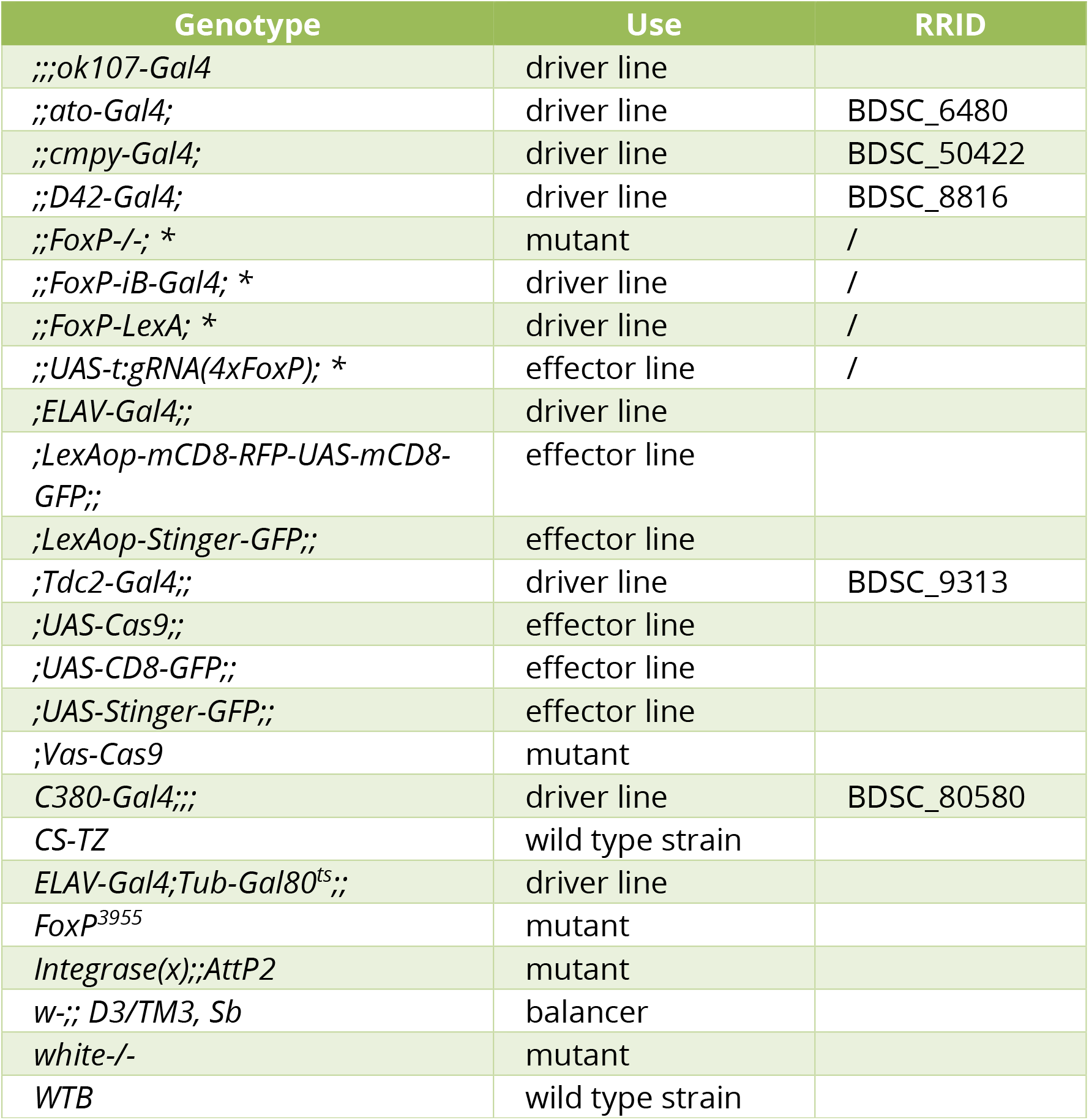
Complete list of the fly lines used in this study. The flies strains marked with “*” are the ones that we created in the present work.

For local knock-out experiments, two genetic constructs need to be brought together for the method to work effectively. The endonuclease Cas9 needs to be present as well as the guide RNA (gRNA) to provide a target for the nuclease. Hence, the appropriate control groups express only one component of the CRISPR/Cas9 combination. One line drives expression only of the Cas9 endonuclease (i.e., *xxx-Gal4>UAS-Cas9,* without gRNAs) and the other drives expression only of the gRNAs (i.e., *xxx-Gal4>UAS-t:gRNA(4×FoxP*) without Cas9). In this fashion, each strain not only controls for potential insertion effects of the transgenes used, but also for potential detrimental effects of expressing the components alone, irrespective of the excision of the target gene.

For the behavioral analysis involving the *FoxP-KO* mutant and the *FoxP-iB-Gal4* driver line we crossed the lines back into *Wild Type Berlin* genetic background for at least six generations in order to get the same genetic background as the *WTB* control.

### 2.2. *In-silico* sequences alignment

The transcript and protein sequences of the different FoxP isoforms were downloaded from https://flybase.org and aligned with *Clus-tal Omega* for multiple sequence alignment. The protein domains were analyzed with the *NCBI Conserved Domain Search* tool, and the stop codons were identified with *ExPASy Translate* tool (Fig. 1A).

### 2.3. Transgenics

We used CRISPR/Cas9 Homology Directed Repair (HDR) to edit the *FoxP* locus (Gratz et al., 2014) and generated t-RNA based vectors for producing multiple clustered regularly-interspaced (CRISPR) gRNAs from a single transcript (Port et al., 2016). We created a total of two driver lines (*FoxP-iB-Gal4* and *FoxP-LexA*), one mutant line (*FoxP-KO*) and one effector line (*UAS-t:gRNA(4×FoxP*)).

#### FoxP-iB-Gal4

To create an isoform-specific driver line, we inserted a Gal4 sequence into exon 8, which is specific to isoform B. Two 1 kb homology fragments were PCR-amplified (primers Hom1: fw 5’-GGGGGCGGCCGCCGTG-GAAGGTAAAATGCCCCATATATG-3’, rv 5’-GGGGCCGCGGCCCTCGTGTAAGGAAAGGTTCG-TACGAATCGC-3’;primers Hom2: fw 5’-GGGGGGCGCGCCACAAGTGCTTTGTAC-GTTATGAA-3’, rv 5’-GGGGGGTACCGGTCAC-TGAGTATCGTTAATGATC-3’) and digested with the appropriate restriction enzymes (Hom1: NotI and SacII, Hom2: AscI and KpnI) to be ligated in the pT-GEM(0) (Addgene plasmid # 62891; RRID:Addgene_62891) vector [25] which contained a Gal4 sequence and a 3xP3-RFP-SV40 sequence for selection of transformants. The gRNA sequences used are: sense 5’-CTTCGACGTACAAAGCACTTGTGTA-3’, and asense 5’-AAACTACACAAGTGCTTTGTACGTC-3’. They were annealed and cloned inside a pU6-gRNA (Addgene plasmid # 53062; RRID:Addgene_53062) vector [26], previously digested with BbsI restriction enzyme.

#### FoxP-LexA

To create a driver line that reflects expression of all *FoxP* isoforms, we inserted a LexA sequence into exon 3. Two 1 kb homology fragments were PCR-amplified (primers Hom1: fw 5’-GGGGGCGGCCGCCAG-GAATGGCGGCATATGAGT-3’, rv 5’-GGGGCCGCGGCCCTCTATTACGGTAAGCG-GACTCCGG-3’; primers Hom2: fw 5’-GGCCGG-TACCATAGCATAGGCCGACCCATC-3’, rv 5’-GGCCACTAGTTCACATTCTCAACCCG-CATAAAGC-3’) and digested with the appropriate restriction enzymes (Hom1: NotI and SacII, Hom2: KpnI and SpeI) to be ligated in the pT-GEM(0) vector which contained a LexA sequence and a 3xP3-RFP-SV40 sequence for selection of transformants. The gRNA sequences used are: sense 5’-CTTCGGGTCGGCCTATGC-TATTTA-3’, asense 5’-AAACTAAA-TAGCATAGGCCGACCC-3’. They were annealed and were cloned inside a pU6-gRNA vector previously digested with BbsI restriction enzyme.

#### FoxP-KO

To prevent expression of any isoform of the *FoxP* gene, we removed part of exon 1, the complete exon 2 and part of exon 3. Two 1 kb homology fragments were PCR-amplified (primers Hom1: fw 5’-GGGGCTAGCCAAAATAA-GATGTGTCTGGTTTCCTTG-3’, rv 5’-GGGCCGCGGGCATGGCGAACTCATCGTG-3’, primers Hom2: fw 5’-GGGGACTAG-TAGAGGGAAAGTTTTGCCGG-3’, rv 5’-GGGGCTGCAG-TATGAAGGGACAGATTGTGCCGG-3’) and digested with the appropriate restriction enzymes (Hom1: NheI and SacII, Hom2: SpeI and PstI) to be ligated in the pHD-DsRed-attP (Addgene plasmid # 51019;RRID:Addgene_51019) vector which contains a 3xP3-DsRed sequence for selection of transformants. The gRNA sequences used are: gRNA1 sense 5’-CTTCGCGGATGA-TAGTACTTCCGCA-3’, asense 5’-AAACTGCG-GAAGTACTATCATCCGC-3’;gRNA2 sense 5’-CTTCGAAGGACGTGCCCGGAAGAGA-3’, asense 5’-AAACTCTCTTCCGGGCACGTCCTTC-3’. They were annealed and were cloned inside a pU6-gRNA vector previously digested with BbsI restriction enzyme.

#### UAS-t:gRNA(4xFoxP)

To create an isoform-unspecific conditional effector line we phosphorylated and annealed 3 sets of oligos (1. fw 5’-CGGCCCGGGTTCGAT-TCCCGGCCGATGCAGAG-CATCGATGAATCCTCAAGTTTCAGAGC-TATGCTGGAAAC-3’, rv 5’-GCTCGGA-TATGAACTCGGGCTGCACCAGCCGG-GAATCGAACC-3’; 2. fw 5’-GCCCGAGTTCATA-TCCGAGCGTTTCAGAGCTATGCTGGAAAC-3’, rv 5’-ACGGCATATGCCATGAGCAATGCAC-CAGCCGGGAATCGAACC-3’; 3. fw 5’-TTGCTCATGGCATATGCCGTGTTTCAGAGC-TATGCTGGAAAC-3’, rv 5’-ATTTTAACTTGC-TATTTCTAGCTCTAAAACAACCATGTTCCG-TATTCAGATGCACCAGCCGGGAATCGAACC-3’) that were cloned with a single Gibson Assembly reaction in a pCFD6 (Addgene plasmid # 73915; RRID: Addgene_73915) vector [27] which was previously digested with BbsI restriction enzyme.

After the constructs were created, they were injected into early embryos (*;Vas/Cas9;* for the *FoxP-iB-Gal4, FoxP-LexA* and *FoxP-KO* and *In-tegrase(x);;AttP2* for *UAS-t:gRNA(4xFoxP)).* The resulting transformants were crossed two times with the balanced flies *w-;; D3/TM3, Sb.*

### 2.4. Immunohistochemistry

Three to six days-old adults were fixated in 4% paraformaldehyde (PFA) at 4 °C for 2 hrs and dissected in 0.01% phosphate-buffered saline with Triton^®^ X-100 detergent (PBST). For larval staining, 3^rd^ instar larvae were selected, dissected in 0.01% PBST and fixated in 4% PFA at room temperature (RT) for 30 min. Clean brains were washed 3 times in 0.01% PBST for a total time of 45 min and then blocked with 10% normal goat serum (NGS) for 1 hr. Subsequently the brains were incubated with the appropriate primary antibody for 1-2 nights at 4 °C (Table 2). After 3 washing steps of 15 min each, the brains were incubated with the secondary antibody (Table 3) for 5-7 hrs at RT. After an additional 15 min washing step, the brains were placed on glass microscope slides and mounted with the antifade mounting medium Vectashield® (Vector Laboratories, Burlingame, CA).

**Table 2:** Complete list of the primary antibodies used in this study

| Antigen | Host | Dilution | Incubation | Source | RRID |
| --- | --- | --- | --- | --- | --- |
| *Chaoptin-11* | mouse | 1:500 | 1 night | DSHB | AB_528161 |
| *ChAT* | mouse | 1:250 | 1 night | DSHB | AB_528122 |
| *ELAV* | rat | 1:100 | 1 night | DSHB | AB_528218 |
| *FoxP* | guinea pig | 1:200 | 2 nights | Lawton et al., 2014 | / |
| *GABA* | rabbit | 1:250 | 2 nights | GeneTex | AB_2037030 |
| *nc82* | mouse | 1:500 | 1 night | DSHB | AB_2314866 |
| *p-SMAD1/5* | rabbit | 1:250 | 2 nights | Cell Signaling Technology | AB_491015 |
| *REPO* | mouse | 1:500 | 1 night | DSHB | AB_528448 |
| *TH* | rabbit | 1:500 | 1 night | Millipore | AB_390204 |

**Table 3:** Complete list of the secondary antibodies used in this study

| Antigen | Host | Dilution | Incubation | Source | RRID |
| --- | --- | --- | --- | --- | --- |
| *Anti-guinea pig Cy<sup>TM</sup>3* | goat | 1:200 | 7 hours | Jackson ImmunoResearch | AB_2337423 |
| *Alexa-Fluor-Anti-mouse 555* | goat | 1:250 | 5 hours | ThermoFisher Scientific | AB_2535844 |
| *Alexa-Fluor-Anti-rabbit 555* | goat | 1:250 | 5 hours | ThermoFisher Scientific | AB_2535849 |
| *Alexa-Fluor-Anti-rat 555* | goat | 1:250 | 5 hours | ThermoFisher Scientific | AB_2535855 |

### 2.5. Image acquisition and analysis

All images were acquired with a Leica SP8 confocal microscope (RRID: SCR_018169), images were scanned at a frame size of 1024×1024 pixels at 200 or 100 Hz. The objectives were 20x dry and 20x/40x/60x oil immersion. Images were processed with ImageJ software (National Institutes of Health, Maryland, USA; RRID: SCR_003070) [28], only general adjustments to color, contrast, and brightness were made. The cell counting was performed with IMARIS 9.0 (Oxford instruments, UK) software on *UAS-Stinger-GFP* stacks, using the tool for spots counting. For the *FoxP-iB-Gal4/FoxP-LexA* count (Fig. 4B), five brains were counted for each genotype at both larval (3^rd^ instar) and adult (2-3 days old) stages. The colocalization analysis was performed with the ImageJ *Colocalization Threshold* tool (Tony Collins and Daniel James White) (Fig 5B). The 3D rendering (Fig. 12B) was performed with IMARIS 9.0 software on the *Drosophila* standard brain from https://www.virtualflybrain.org.

### 2.6. RT-qPCR

The knockout efficiency was assessed using RT-qPCR (see Figs. 1D, 4E). We extracted RNA from 20 flies for each genotype (white^-^, heterozygous mutant and homozygous mutant, both with a white^-^ background), following the TriFast™ protocol from peqlab (a VWR company) (Catalog # 30-2010). The RNA was subsequently transcribed into cDNA using the OneStep RT-PCR Kit from QIAGEN (Catalog # 210212) with the following thermocycler program: 42 °C for 2 minutes, 4 °C pause until manual restart at 42 °C for 30 minutes, 95 °C for 3 minutes and finally 10 °C ∞. Subsequently we performed the qPCR. Primer sequences were identical to those used by [18]. For the qPCR reaction we used a Bio-Rad CFX Connect Real-Time PCR Detection System thermocycler and the Bio-Rad CFX manager software to store and analyze the data. Every sample was run in triplicate in a 96-well plate in a total volume of 10 μl. The mixture contained 5 μl sybrGreen master mix (ORA™ qPCR Green ROX H Mix, 2X, from highQu, Catalog # QPD0201), 0.5 μl from each primer, 1 μl of 1:10 diluted cDNA and 3 μl H_2_O. As reference, we used the housekeeping gene *rp49 (ribosomal protein 49),* while as a negative control we used the same reaction mix without cDNA. The qPCR reaction program used was: 95 °C for 2 minutes, 95 °C for 10 seconds, 60 °C for 10 seconds, 65 °C for 30 seconds (from step 2 to 4 × 39 rounds), 95 °C for 10 seconds, and finally from 65 °C to 95 °C for + 0.5 °C / 5 s. The experiments were repeated 2 to 4 times.

### 2.7. Behavior

All behavioral experiments were performed in Buridan’s paradigm (RRID: SCR_006331) [29]. In this experiment, we analyzed both temporal components of walking behavior (often subsumed under ‘general locomotion’) and spatial components such as fixation of landmarks or the straightness of the walking trajectory. Buridan’s paradigm (Fig. 6A) consists of a round platform with a diameter of 117 mm which is surrounded by a water-filled moat. The platform is situated at the bottom of a uniformly illuminated white cylinder, 313 mm in height and 293 mm in diameter [30]. Two black stripes are placed on the inside of the cylinder, opposite each other, serving as the only visual cues for the flies. Two days-old female flies were collected and their wings were clipped under CO_2_ anesthesia. After one night recovery at 25°C they were tested in Buridan’s paradigm for 15 minutes (doi: 10.17504/protocols.io.c7vzn5). The position of the fly is recorded by a camera (Logitech Quickcam Pro 9000) connected to a computer running our BuriTrack software (http://buridan.source-forge.net)

The analysis software CeTrAn [30] (https://github.com/jcolomb/CeTrAn) extracts a variety of parameters from the scored trajectories. From the parameters extracted by CeTrAn, we used the temporal parameters median speed (the median of all instantaneous speed data points measured when the fly is walking), distance travelled, number of walks (sections of the trajectory which connect the platform areas closest to the two stripes) and activity time (fraction of time spent walking), as well as the spatial parameters stripe deviation (angular deviation of the fly’s heading from the center of the stripe in the frontal visual field) and meander (a measure of the tortuosity of the fly’s trajectory). Transition plots visualize the areas on the platform that the flies most frequently visited. More details in [30].

For the experiment involving *Tub-Gal80^ts^* (Fig. 8E-H; 11A-B), flies were raised at 18 °C, moved to 30 °C for 12 hrs (embryos) or 48 hrs (pupae and adults) and subsequently left at 25 °C for the rest of the development (embryos and pupae) or overnight for recovery (adults) before testing.

### 2.8. Statistical analysis

All graphs were created and statistical analysis was performed using GraphPad Prism 6 (GraphPad Software, Inc., California, USA; RRID: SCR_002798) software. Sample variances were compared with an F-test. In the absence of significantly different variances, we used Student’s t-tests (two-tailed) or one way ANOVAs followed by Tukey’s post hoc test for multiple comparisons. If the F-test was significant at p<0.005 (see below), we used a Mann-Whitney U-test or a Kruskal-Wallis ANOVA followed by Dunn’s post hoc test for multiple comparisons. Alpha values were set to 0.5%, in order to reduce the chances of false-positives, following the arguments detailed in [31], where BB is an author.Whenever null hypothesis significance testing was performed using the nonparametric tests, it is indicated in the figure legends. All other tests were performed using parametric tests.

The initial behavioral experiments (Fig. 6) were carried out with a sample size which, from experience, would be sufficient to detect medium to large effects, i.e., N~20. We then used these results to perform a power analysis for the subsequent experiments. We found that effect sizes such as those exhibited in the speed, meander or stripe fixation parameters required a sample size of up to 18 to reach 80% statistical power at an alpha of 0.5% [31], while effects such as those in the activity time parameter would require up to 100 flies. We corroborated these analyses with Bayesian analyses, where the activity time parameter yielded a Bayes factor below one, while the other effects yielded Bayes factor values beyond 100. Therefore, we set the target sample size for all subsequent Buridan experiments to 18 and p<0.005 was considered significant. Data are expressed as averages ± SEM or averages ± SD, and each case is indicated in the legend of each figure.

### 2.9. Data deposition

All raw data is publicly accessible with an Attribution 4.0 International (CC BY 4.0) license. Data sources for specific figures are referenced in the text. A compressed file with all the data is also available [32].

## 3. Results

### 3.1. *FoxP-iB* expression in the *Drosophila* brain

The *FoxP* transcription factor binds DNA with the forkhead (FH) box domain (Fig. 1A, yellow boxes;raw data deposited at doi: 10.6084/m9.figshare.12607700). The gene consists of 7 introns and 8 exons. The FH box is split into different segments spanning exons 6, 7 and 8. The last two exons (7 and 8) are subjected to alternative splicing, leading to two different protein isoforms: isoform A (*FoxP-iA),* which results from splicing exon 6 to exon 7, and isoform B (*FoxP-iB*) where instead exon 8 is spliced to exon 6. A third, intron-retention isoform is transcribed by failing to splice the intron between exon 6 and 7 out (*FoxP-iIR).* While the first two isoforms contain a complete and putatively functioning FH box (with 10 amino acids different between the two; Fig.1A, dashed box), the putative FoxP-iIR FH box appears to be truncated due to a stop codon in the intron sequence (Fig. 1A, red line), putting the transcription factor function of this isoform in doubt. Of the three FoxP isoforms, *FoxP-iB* was most directly associated with the learning phenotype discovered by [18]. Therefore, we inserted the sequence of the yeast transcription factor Gal4 into exon 8, the exon which is exclusive to *FoxP-iB* (Fig. 1A). This insertion leads to the expression of the Gal4 transcription factor only in *FoxP-iB* positive cells. At the same time, the insertion also disrupts the FH box DNA binding domain of the *FoxP* gene, putatively preventing the FoxP protein to act as a transcription factor, effectively mutating the gene for this function.

Observing Gal4 expression with different green fluorescent proteins (GFPs) under control of the UAS promoter (to which Gal4 binds), revealed that *FoxP-iB* is expressed throughout the whole development of the fly, from embryo (data deposited at doi: 10.6084/m9.figshare.12607652) to adult, in both brain and ventral nerve cord (VNC) (Fig. 1B). In 3^rd^ instar larvae, we observe expression in the central brain (but not in the optic lobes) and in the anterior portion of the VNC. In the adult nervous system, the main neuropil expression domains comprise protocerebral bridge, gnathal ganglia (subesophageal zone), vest, saddle, noduli, and superior medial protocerebrum. GFP-positive cell body clusters could be found in the cortex of the central brain and around the optic lobes (Fig. 1B).

We next validated the expression pattern of our *iB*-specific driver line to the staining of an available isoform unspecific polyclonal antibody [21]. We observed complete colocalization of the driver line with the antibody staining in both larvae and adults, i.e., there were no GFP-positive cells that were not also labeled by the FoxP antibody (Fig. 1C). The cells only stained for the FoxP antibody and not for GFP are presumably cells expressing the other *FoxP* isoforms (*FoxP-iA* and *FoxP-iIR,* Fig. 4). Notably, in contrast to previous reports [33,34] but consistent with [20], we did not detect any *FoxP* expression in mushroom body neurons, neither with our driver line, nor with the antibody.

Postulating that our transgene disrupted expression of the *FoxP* gene, we measured mRNA levels of all three isoforms with RT-qPCR (Fig. 1D). With one of the primers placed over the Gal4 insertion site, we observed approximately half the wild type *FoxP-iB* expression levels in heterozygous animals, while wild type *FoxP-iB* expression was essentially abolished in the homozygous transgenes. We did not observe any significant changes in the other two isoforms. However, this disruption of the *FoxP-iB* isoform did not have an effect on the development of the *FoxP*-positive neurons, as both heterozygous and homozygous *FoxP-iB-Gal4* mutants showed the same number of GFP-labelled cell nuclei (data deposited at doi: 10.6084/m9.figshare.12932705).

### 3.2. *FoxP-iB* is expressed in a variety of neurons, but not glia

With *FoxP* involved in learning and expression patterns suggesting neuronal expression (Fig. 1), we investigated whether the observed expression was exclusively neuronal, or if there were also *FoxP-iB* expressing glial cells. Therefore, we stained 3^rd^ instar larva and adult brains with antibodies against ELAV (neuronal marker) and REPO (glial marker). At both developmental stages, the two stainings reveal exclusive *FoxP-iB-*mediated GFP colocalization with ELAV without any colocalization with REPO (Fig. 2A-B; raw data deposited at doi: 10.6084/m9.figshare.12607706), suggesting that *FoxP-iB* is expressed exclusively in neurons. These data are consistent with results published previously [20,35], validating the methods employed here.

**Fig. 2:**
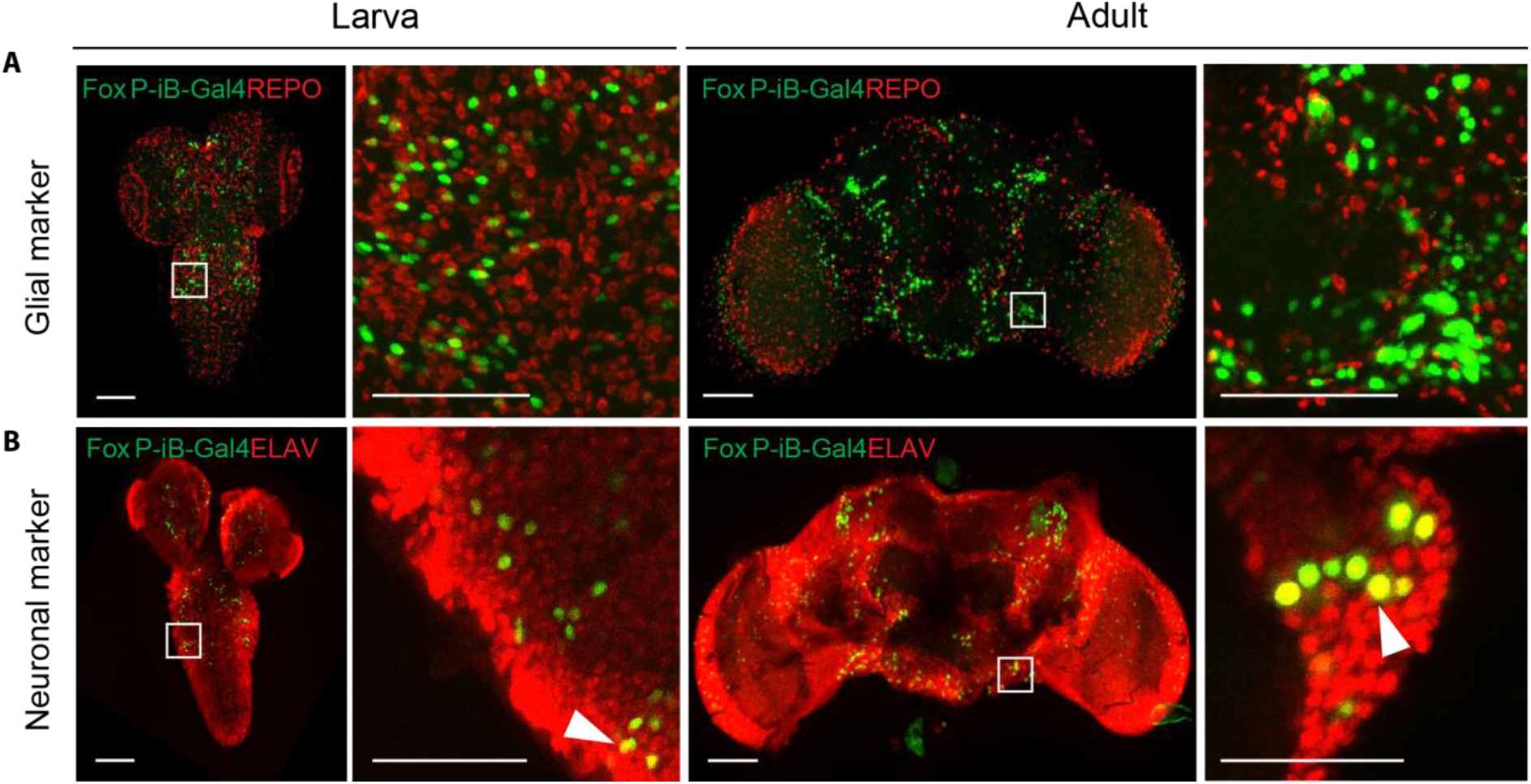
*Only neurons, not glia, are expressing* FoxP-iB *in the* Drosophila *brain.* Immunohistochemistry on *FoxP-iB-Gal4>Stinger-GFP* flies with REPO (glia, A) and ELAV (neurons, B) markers. Note the lack of colocalization of *FoxP-iB* driven GFP with the glial marker in both 3rd instar larvae and adult brains (A). In contrast, exclusive colocalization of *FoxP*-driven GFP with the neuronal marker was observed in both developmental stages (white arrowheads indicate typical examples). Scale bars: 50 μm.

We next investigated in more detail the types of neurons in which *FoxP-iB* is expressed. Using a variety of antibodies (Fig. 3; raw data deposited at 10.6084/m9.figshare.12607712) used as markers for different neuronal cell types we detected *FoxP-iB* expression in most of the cell types investigated. For technical reasons, we stained adult brains only with the anti-TH antibody, while the remaining markers were used on larval nervous systems. Extensive colocalization was observed with p-SMAD1/5 (a motorneuron marker) in the VNC but not in the central brain (CB) (Fig. 3A). Some *FoxP-iB* neurons were positive for ChAT (cholinergic) or GABA (inhibitory) both in the VNC and in the CB (Fig. 3B-C). Finally, a few *FoxP-iB* positive neurons were found to colocalize with Tyrosine hydroxylase (dopaminergic neurons) in the CB only (Fig. 3D). These data are consistent with the study performed by *[36]* in honeybees where they found colocalization between Am-FoxP positive neurons and GABAergic, cholinergic and monoaminergic markers. No colocalization was instead found in photoreceptor cells stained with Chaoptin (data deposited at doi: 10.6084/m9.figshare.12607643), a marker for photoreceptor cells [37].

**Fig. 3:**
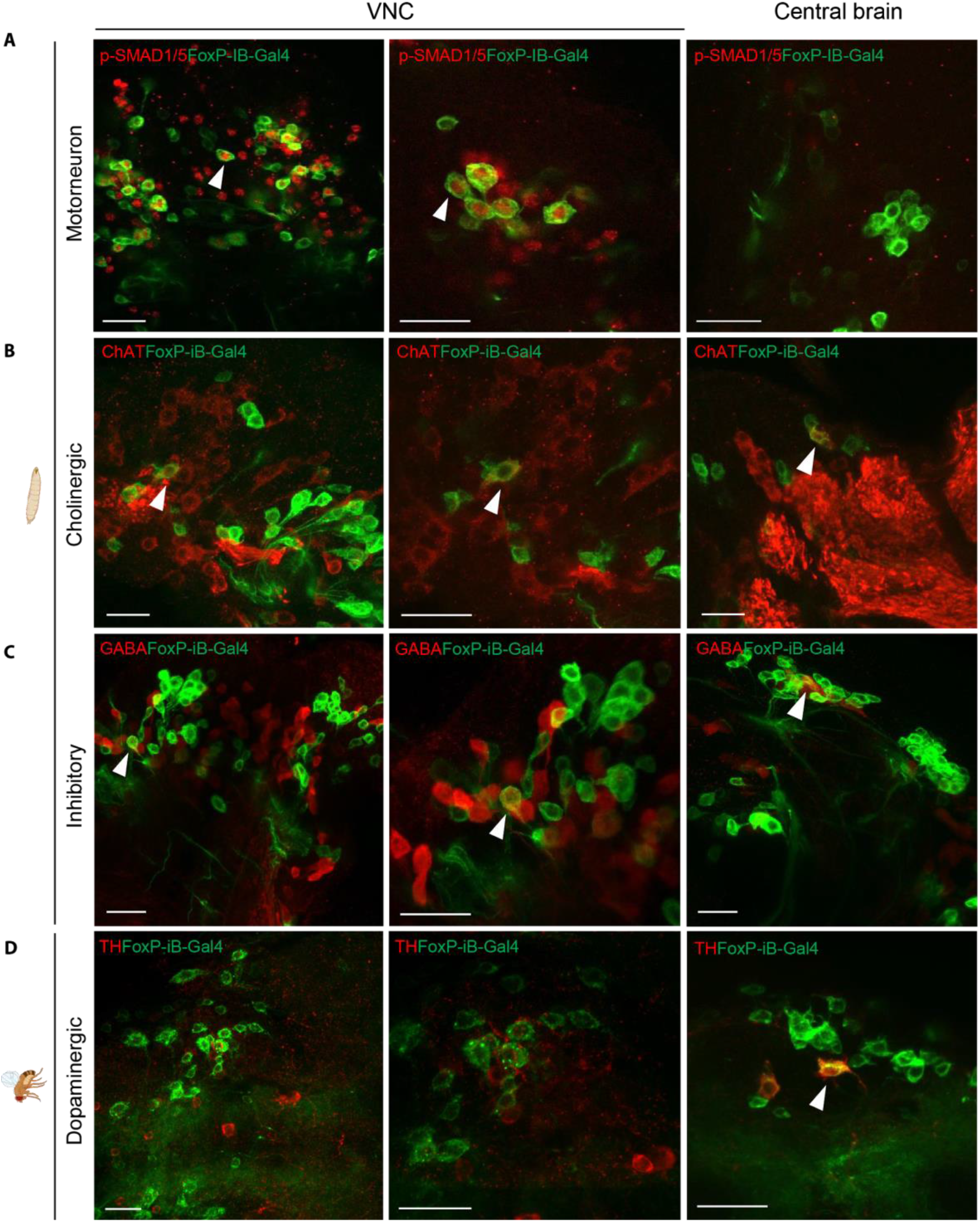
FoxP-iB *is expressed in various types of neurons.* Immunohistochemistry on *FoxP-iB-Gal4>CD8-GFP* larvae and adults using different antibodies. (A) Some of the *FoxP-iB* positive neurons colocalize with p-SMAD1/5 in the VNC but not in the central brain. (B-C) *FoxP-iB* neurons positive for ChAT or GABA have been found in both the VNC and CB. (D) Only few *FoxP-iB* neurons colocalize with TH and only in the CB. White arrowheads indicate examples of colocalization. Scale bars 25 μm. (A-C): larvae; (D): adult flies.

### 3.3. *FoxP-iB* is expressed in a subset of *FoxP-ex*-pressing neurons

As the antibody staining against the FoxP protein indicated more cells expressing FoxP than our *FoxP-iB*-specific driver line was reporting (Fig. 1B), we created a second driver line, designed to drive expression in all FoxP cells, irrespective of isoform. We inserted a sequence for the bacterial LexA [38] transcription factor into exon 3 (Fig. 4A; raw data deposited at doi: 10.6084/m9.figshare.12607730). Comparing *Stinger-GFP* expression from each driver line revealed a more expansive pattern for the isoform-unspecific driver (Fig. 4B), one that matches very well with the FoxP antibody staining (Fig. 1B). This visual impression was corroborated by a quantification of stained nuclei comparing between both drivers (Fig. 4C). This quantification allowed us to trace the proliferation of FoxP cells from around 500 in 3^rd^ instar larvae to around 1800 in three days-old adults. In contrast, there are only about 300 cells expressing *FoxP-iB* in the 3^rd^ larval instar and only around 1300 in three days-old adults. We noticed that the largest differences in terms of cell number between *FoxP-LexA* and *FoxP-iB-Gal4* flies (both larvae and adults) were found in the CB, while the VNC numbers differed considerably less. Taken together, in 3^rd^ instar larvae and in three days-old adults, 66% and 65%, respectively, of the total number of *FoxP* neurons in the *Drosophila* nervous system express *FoxP-iB.As* with our previous insertion, also this one was expected to disrupt expression of the *FoxP* gene. To investigate the extent of this disruption on the mRNA level, we again performed RT-qPCR. As expected, this insertion affected all isoforms. In heterozygous flies, the expression level was increased, while in homozygous flies it was decreased (Fig. 4E). It is important to note that here, in contrast to the *FoxP-iB* insertion, all primers are binding sequences downstream of the insertion site (see Fig. 1A, D) and so are amplifying all transcripts, with or without the insertion. Possibly, the FoxP protein is involved in its own regulation such that in heterozygotes, the missing gene copy leads to a compensatory increase in transcription rate, but in homozygous mutant animals without any FoxP protein, only basal expression levels remain.

**Fig. 4:**
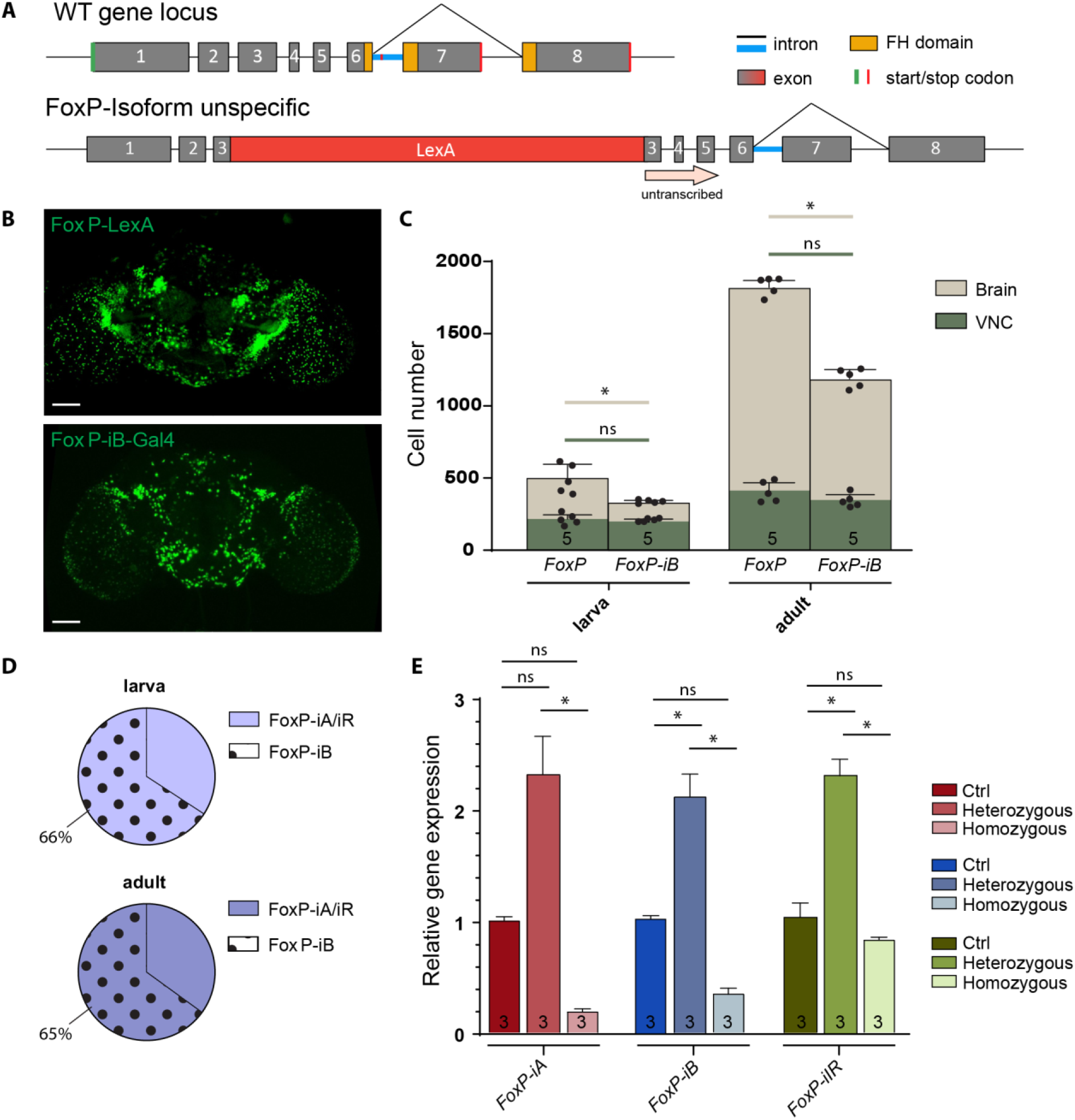
FoxP-iB *is expressed in a subset of FoxP-expressing neurons*. (A) Schematic representation of the FoxP gene locus after LexA insertion. This is an isoform unspecific construct with the insertion of a LexA sequence in exon 3. (B) Expression pattern of *FoxP-LexA* and *FoxP-iB-Gal4* driving *Stinger-GFP.* (C) Cell counting performed with IMARIS on *FoxP-LexA* and *FoxP-iB-Gal4>Stinger-GFP* (3rd instar larvae and 3 days-old adults) in both brain and VNC. (D) Pie charts that summarize the results from (C). (E) RT-qPCR on FoxP-LexA flies (control, hetero- and homozygous flies). Data are expressed as means ± SD in (C) and as means ± SEM in (E). *p<0.005. Scale bars: 50 μm.

In order to directly compare the expression patterns of our two driver lines, we used them to drive reporter genes fluorescing at different wavelengths (i.e., *LexAop-RFP;UAS-CD8-GFP*) and analyzed their patterns in adults (Fig. 5; raw data deposited at doi: 10.6084/m9.figshare.12607763). In this way, we labeled all *FoxP*-expressing neurons red and neurons that specifically expressed *FoxP-iB* green. We used the “Colocalization Threshold” tool from ImageJ, which computes false colors to enhance the comparison between the two driver lines and let the differences stand out (see M&M). The other two isoforms are expressed in additional areas (the antennal lobe, the lobula, the fan shaped body and the posterior ventrolateral protocerebrum) compared to *FoxP-iB* expression (Fig. 5A, red arrowheads; Fig. 5B, blue areas).

**Fig. 5:**
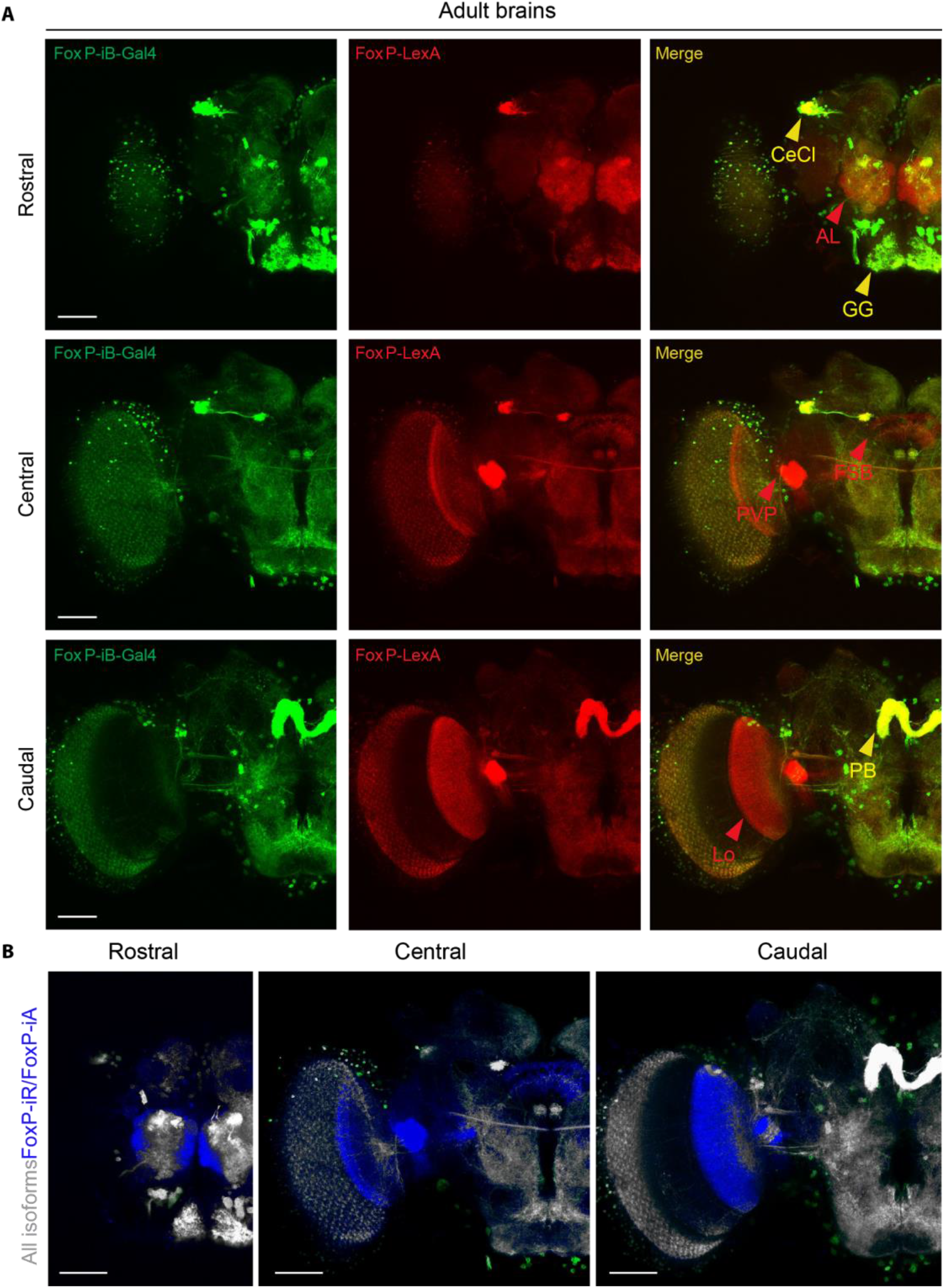
*Comparison of* FoxP *isoform expression patterns.* (A) Confocal images of adult brains expressing *FoxP-iB-Gal4>CD8-GFP* (green) and *FoxP-LexA>CD8-RFP* (red). (B) Difference rendering (blue) indicating areas expressing only non-iB isoforms. AL: antennal lobe, PVP: posterior ventrolateral protocerebrum, FSB: fan shaped body, Lo: lobula, cecl: cell cluster, GG: gnathal ganglion, PB: protocerebral bridge). Scale bars: 50 μm.

We also crossed this *FoxP-LexA* line with *LexAop-RFP-UAS-CD8-GFP* and *Tdc2-Gal4* to investigate any potential tyraminergic or octopa-minergic *FoxP* neurons, but despite a close proximity between the two cell types, no colocalization was found (data deposited at doi: 10.6084/m9.figshare.12607670).

### 3.4. *FoxP-iB* knockout flies show a multitude of behavioral abnormalities

Mutations in the *FoxP* gene do not only affect operant self-learning, for instance, different alleles also affect flight performance and other locomotion behaviors to different degrees [18,20,21]. Because of the *FoxP* pleiotropy affecting various innate motor behaviors independently from motor learning, we turned to Buridan’s paradigm [29,30] as a powerful tool to measure several locomotor variables. Buridan’s paradigm allows us to test a broad panel of behavioral parameters covering temporal parameters such as speed or general activity time and spatial parameters such as the straightness of a fly’s trajectory (meander) or the degree to which the animal is heading towards one of the two vertical landmarks (stripe fixation; Fig. 6A).

**Fig. 6:**
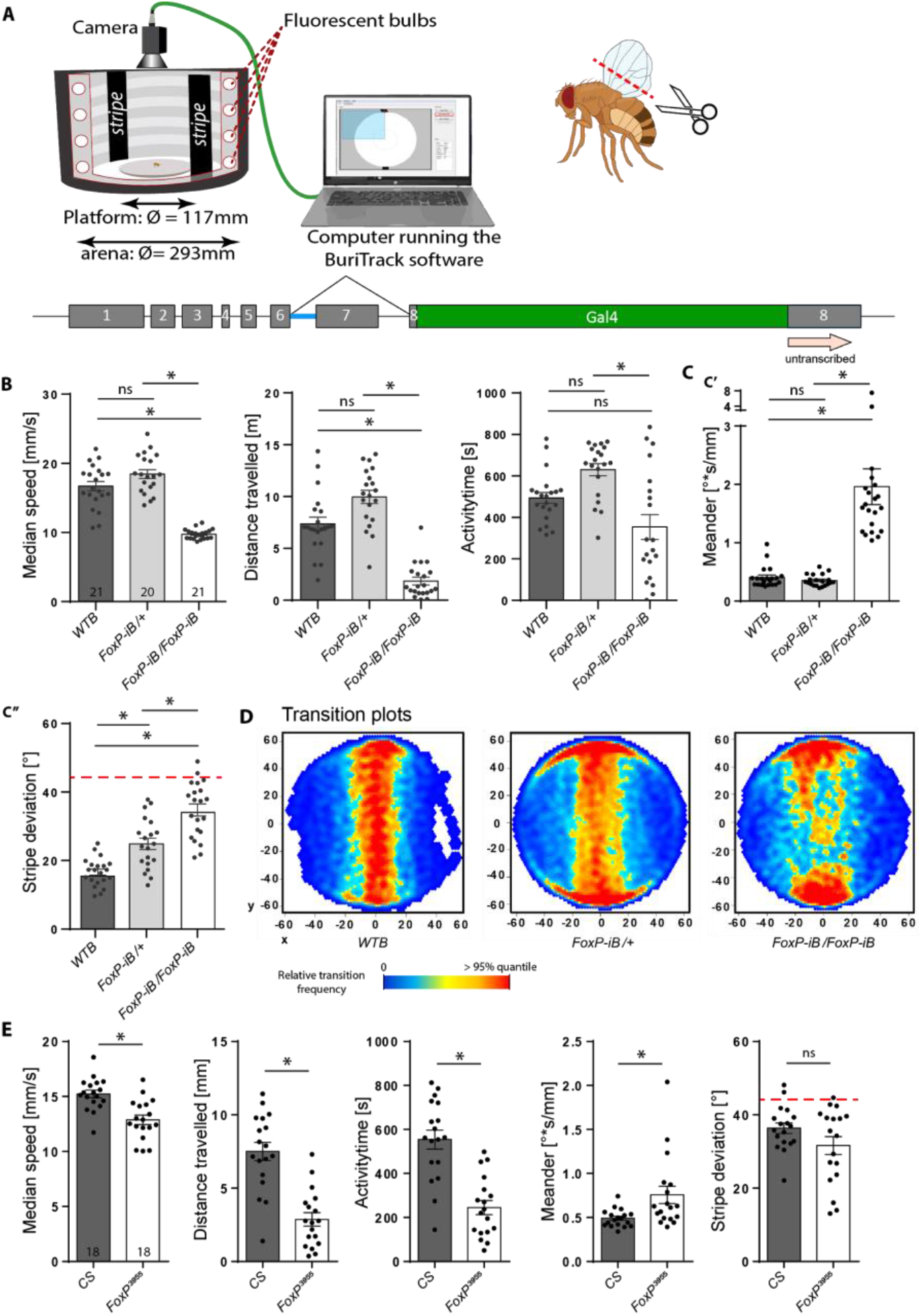
FoxP-iB *mutant flies are impaired in several parameters in Buridan’s paradigm.* (A) Schematic of Buridan’s paradigm. A fly with shortened wings is put in the center of a platform inside a circular arena with two opposing black stripes on the walls. A camera records the position of the fly and the BuriTrack software stores the position data for later analysis with CeTrAn. (B) Temporal parameters. Median speed denotes the instantaneous speed when a fly is walking. Activity time denotes the time spent walking. Distance traveled measures the distance covered by the fly during the experiment. Nonparametric tests for walking speed and activity time. (C) Spatial parameters. Meander is a measure for the tortuosity of a fly’s trajectory. Stripe deviation measures the angular deviation from heading towards the center of the stripe to which the fly is oriented. Red dashed line indicates angular stripe deviation of a random walk. Nonparametric test for meander. (D) The transition plots show the distribution of the platform locations that the flies transitioned through. (E) Temporal and spatial parameters from CS and *FoxP^3955^* mutant flies in Buridan’s paradigm are largely consistent with those from the *FoxP-iB* knock-out. Nonparametric test for meander. *p<0.005.

With our insertions constituting novel alleles impairing *FoxP* expression (Figs. 1, 4), we started by testing the heterozygous and homozygous driver strains without any effectors.

Consistent with previous findings of impaired locomotor behavior in *FoxP* manipulated flies [18,20,21] and the qPCR results showing reduced *FoxP* expression (Fig. 1D), our *FoxP-iB-Gal4* insertion shows abnormalities in Buridan’s paradigm both in temporal and in spatial parameters (Fig. 6; raw data deposited at doi: 10.6084/m9.figshare.12607769). While the homozygous flies walked more slowly, spent more time at rest and fixated the stripes less strongly than wild type control flies, heterozygous flies only differed from wild type flies in their stripe fixation (Fig. 6C). With different effect sizes in each parameter, we selected two representative parameters each for the temporal and the spatial domain, respectively, for comparison of all subsequent lines: walking speed, activity time, meander and stripe fixation.

Because our insertion is located in the same exon as the insertion in the *FoxP^3955^* mutant, we tested the *FoxP^3955^* mutant flies in Buridan’s paradigm and found changes in several temporal parameters, similar to those observed in our driver line (Fig. 6D). However, for the spatial parameters (stripe deviation and meander) only stripe fixation appears normal in these flies, the increased meander indicates that the flies have problems walking straight, despite clearly walking towards the stripes. Thus, in addition to the deficits in operant self-learning and flight performance as reported previously [18], the *FoxP^3955^* mutant flies are also deficient in several temporal and one spatial parameter of walking behavior in Buridan’s paradigm. This walking phenotype is consistent with previous findings of walking deficits associated with *FoxP* manipulations [20,21], but was not detected in a previous publication where tests for walking deficits had been performed [34].

### 3.5. Knocking out all *FoxP* isoforms has similar effects as *FoxP-iB* knockout

With such dramatic motor alterations when only *FoxP-iB,* which is expressed in about 65% of all *FoxP*-positive neurons (Fig. 4), is removed (Fig. 6), it is interesting to study the effects of removing the remaining isoforms for a complete *FoxP* knockout. To avoid unwanted potential side-effects of expressing a different protein in its stead, we created a third fly line where the entire second exon is removed together with parts of exons 1 and 3. We validated this mutant with the polyclonal antibody we used before (Fig. 1). While the antibody detected the FoxP gene product in control flies, there was no signal in our homozygous knockout flies (Fig. 7A, B; raw data deposited at doi: 10.6084/m9.figshare.12607796).

**Fig. 7:**
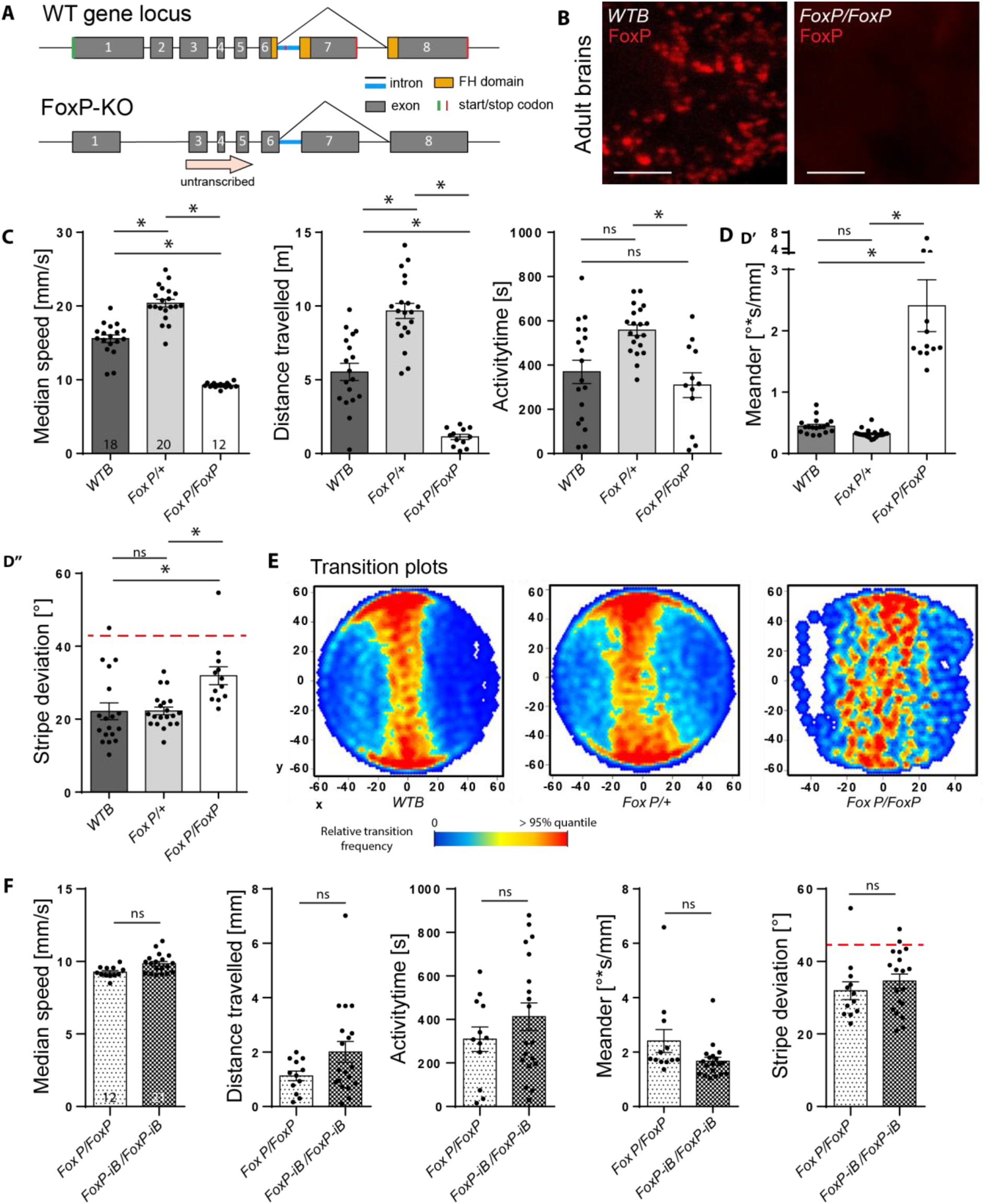
*Deleting the entire* FoxP *gene has similar consequences in Buridan’s paradigm as deleting only* FoxP-iB. (A) Schematic representation of the deletion (*FoxP-KO*) and the wild type (*WT*) gene locus. (B) Immunohisto-chemistry staining for the FoxP gene product in wild type and FoxP-KO mutant brains. (C) Temporal parameters. See Fig. 6 and M&M for definitions. Note the overdominance of the heterozygous *FoxP-KO* flies. Stripe deviation (D) and transition plots (E) show weaker stripe fixation of homozygous FoxP-KO flies. (F) Comparing *FoxP-KO* and *FoxP-iB* flies reveal only a small difference in walking speed. Nonparametric tests for distance travelled and meander. *p<0.005. Scale bars: 25 μm.

Analogous to the behavioral characterization in the *FoxP-iB* insertion line, we tested both heterozygous and homozygous *FoxP-KO* deletion mutants in Buridan’s paradigm (Fig. 7C-F). The results of this experiment closely resemble the ones from the *FoxP-iB-Gal4* insertion line, with homozygous mutants being both significantly less active (Fig. 7C) and fixating the stripes less strongly than the heterozygous mutants and the wild type controls (Fig. 7D, E). For this allele, the heterozygous *FoxP-KO* mutants show higher values for all temporal parameters compared to the wild type controls, while there is no difference in stripe deviation. This trend was already observed in *FoxP-iB* mutants but failed to reach statistical significance. Thus, the *FoxP-KO* allele exhibits differential dominance, with recessive modes of inheritance in some traits and, e.g., overdominance in others. A direct comparison of the data from the two homozygous alleles (*FoxP-iB* and *FoxP-KO*) showed no significant differences (Fig. 7F). Thus, removing the other *FoxP* isoforms had few effects beyond the consequences of removing only *FoxP-iB* alone.

### 3.6. Local and conditional *FoxP* knock-out (*FoxP-cKO*)

Given the patchy expression pattern of *FoxP* in the fly’s nervous system (Figs. 1–5) and the grave consequences for behavior in Buridan’s paradigm if it is manipulated (Figs. 6, 7), we sought to investigate when and where *FoxP* is required for normal walking behavior. To this end, we designed a fourth fly strain which carries a UAS-controlled effector (Fig. 8A; raw data deposited at doi: 10.6084/m9.figshare.12607805). The four guide RNAs (gRNA) each target a different section of the *FoxP* gene (see M&M). If expressed together with the endonuclease Cas9, this effector efficiently mutates the targeted gene [27,39]. We validated this approach by driving both our gRNAs as well as Cas9 using the panneuronal *elav-Gal4* driver and monitoring *FoxP* expression with the FoxP antibody used before (Fig. 8B). Flies with this pan-neuronal excision of the *FoxP* gene (*FoxP-cKO*) were also tested in Buridan’s paradigm and showed even more severe impairments than flies homozygous for a constitutive deletion of the gene (Fig. 8C). In fact, the mutated flies walked so little, that analysis of spatial parameters was not meaningful (Fig. 8D). To allow for temporal control of transgene expression, we also validated the use of the temperature-sensitive suppressor of Gal4, Gal80^ts^ (Fig. 8E). The constitutively expressed Gal80^ts^ prevents Gal4 from activating transcription of the UAS-controlled transgenes until the temperature is shifted from 18 °C to 30 °C, at which point the repressor becomes inactivated and Gal4-mediated transcription commences [23,24]. Using this system to drive gRNA/Cas9 expression for 12 hours in the embryo phenocopies both the mutant and the local phenotypes not only on the protein (Fig. 8F), but also on the behavioral level (Fig. 8G, H). In both experiments, the effects of the manipulations were so severe, that it was not possible to reach the target sample size of 18.

**Fig. 8:**
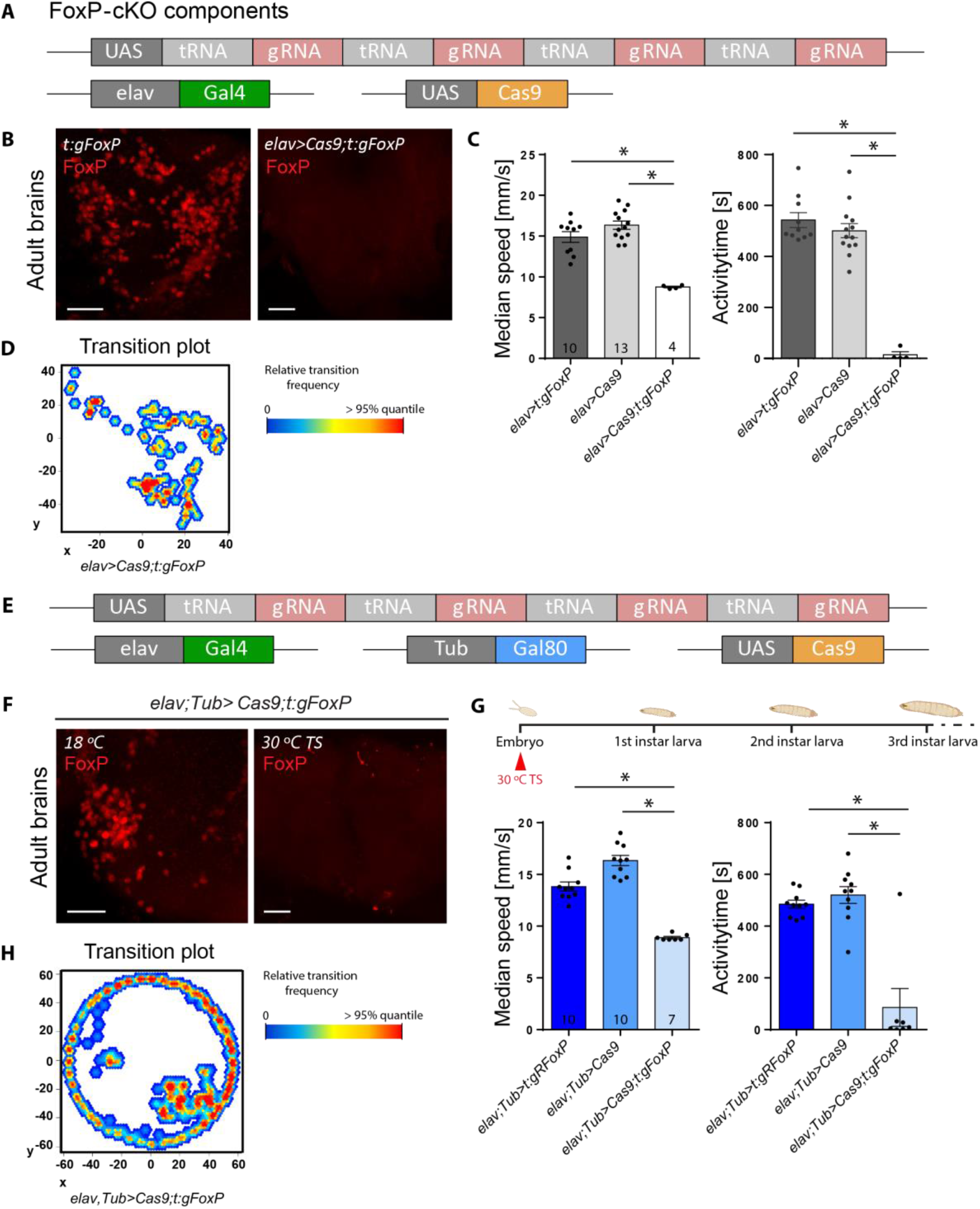
*Local and conditional* FoxP *gene knock-out mimics mutant phenotype.* (A) Construct schematic of the effector (UAS) line we created, together with the *elav* driver line and *Cas9* effector. (B) Immunohistochemistry on adult brains of the effector control line (left), and of the experimental cross (right). Driving expression of our gRNA construct with *elav-Gal4* leads to a highly efficient *FoxP* gene knock-out. (C) This knock-out is also validated in Buridan’s paradigm, where the experimental flies show strongly reduced locomotor activity. Nonparametric test for speed. (D) Transition plot showing the reduced activity of *FoxP-cKO* flies. (E) Schematic of the used transgenic elements. Gal80^48ts^ inhibits Gal4 under 30 °C. (F) Induction of panneural gRNA expression in the embryo eliminates FoxP expression as tested with a FoxP antibody (right) compared to uninduced controls (left). (G-H) Inducing panneural gRNA expression in the embryo also leads to similar locomotor defects as observed in mutants and in flies expressing the gRNAs without temporal control. Nonparametric tests for both parameters. *p<0.005. Scale bars: 25 μm

### 3.7. Local *FoxP-KO:* brain regions and neuron types

Recently, Linneweber *et al*. [40] described the consequences of silencing dorsal cluster neurons (DCNs) on stripe fixation behavior in Buridan’s paradigm. The *FoxP-iB* expression pattern suggests that at least some of these DCNs express FoxP (Fig. 1). Comparing our isoform-unspecific *FoxP-LexA* expression pattern with that of the *atonal-Gal4* line used to drive expression in DCNs [41], we observed substantial overlap (Fig. 9A; raw data deposited at doi: 10.6084/m9.figshare.12607811). Therefore, we used *ato-Gal4* to excise the *FoxP* gene specifically in DCNs. Remarkably, this manipulation did not have any effect on the flies’ behavior in Buridan’s paradigm (Fig. 9B).

**Fig. 9:**
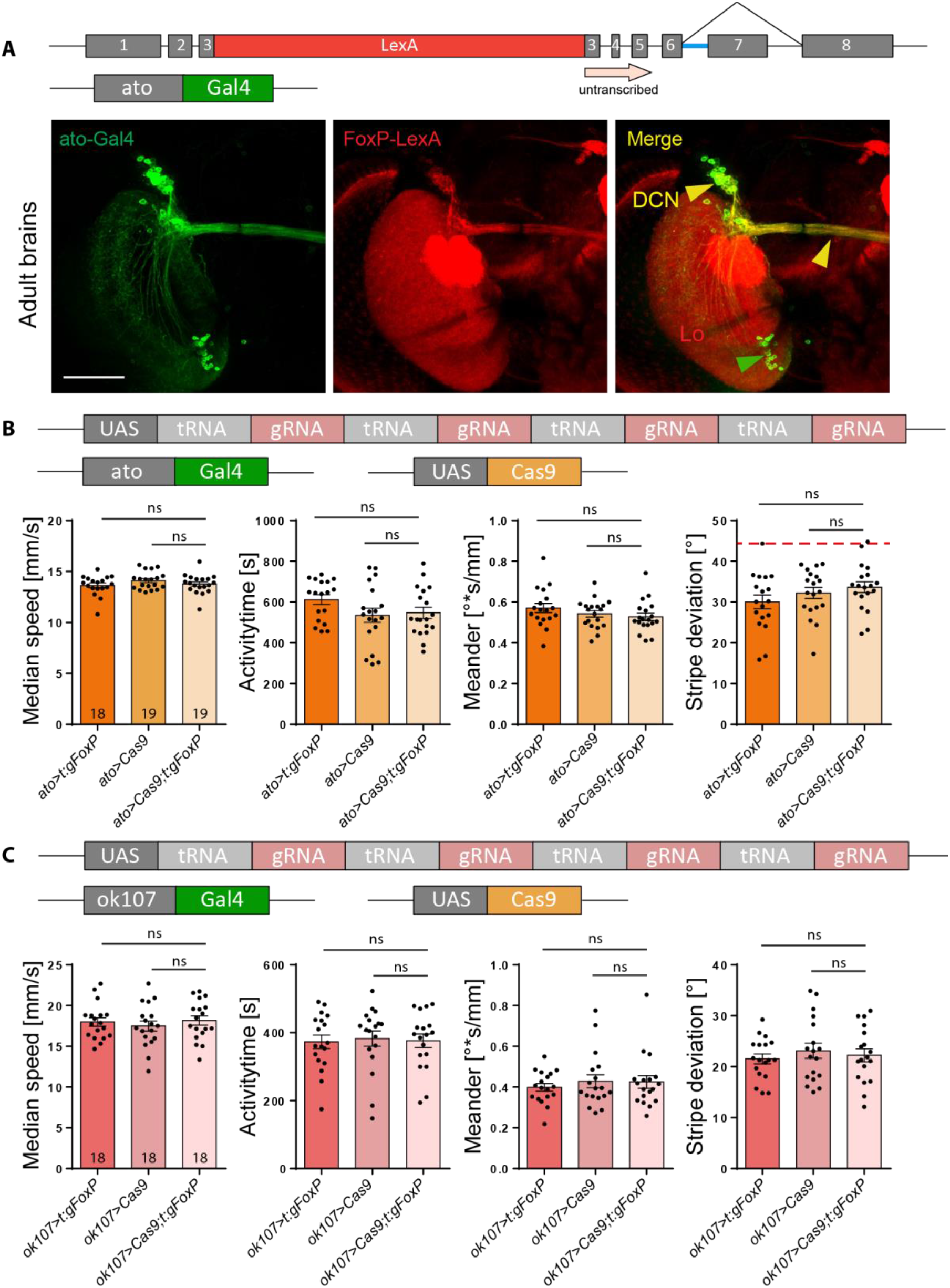
*Local* FoxP-KO *shows no effect in dorsal cluster neurons or mushroom bodies.* (A) Immunohistochemistry showing *FoxP-LexA* expression in the adult brain compared to the expression of the *ato-Gal4* driver. DCN: dorsal cluster neurons, Lo: lobula, Scale bars: 50 μm. (B) Temporal and spatial parameters from Buridan’s experiment show no effects of knocking out the *FoxP* gene in dorsal cluster neurons using the *ato-Gal4* driver. (C) Knocking out the *FoxP* gene in the mushroom bodies using the *ok107-Gal4* driver has no effect on either spatial or temporal parameters in Buridan’s paradigm.

The insect mushroom-bodies (MBs) are not only known as a center for olfactory learning and memory [e.g., 42,43-49], they are also involved in the temporal and spatial control of locomotor activity [e.g., 50,51-59]. In addition, [20] found a subtle structural phenotype in a sub-section of the MBs without detectable *FoxP* expression in the MB Kenyon cells themselves. Finally, there are two reports that expressing anti-*FoxP* RNAi constructs exclusively in the MBs can have behavioral effects [33,34]. For these reasons, despite neither [20] nor us being able to detect any FoxP expression in the MBs (and current RNAi constructs fail to knock down *FoxP* mRNA, see below), we deleted the *FoxP* gene from MB Kenyon cells using the *ok107-Gal4* driver and tested the flies in Buridan’s paradigm. We did not detect any differences to control flies in these experiments (Fig. 9C).

There are two reasons for knocking out *FoxP* in motorneurons, besides *FoxP* expression there (Fig. 3): first, networks of motorneurons in the VNC control movement patterns and walking is directly affected by our manipulations so far (Figs. 6, 7). Second, motorneurons were shown to be important for the type of operant self-learning that also requires *FoxP* [60]. Driving expression of gRNA/Cas9 with either of two motorneuron-specific driver lines (*D42-Gal4* and *C380-Gal4*) led to a significant alteration in locomotor activity in Buridan’s paradigm, both for spatial and for temporal parameters. (Fig. 10A, B; raw data deposited at doi: 10.6084/m9.figshare.12607823).

**Fig. 10:**
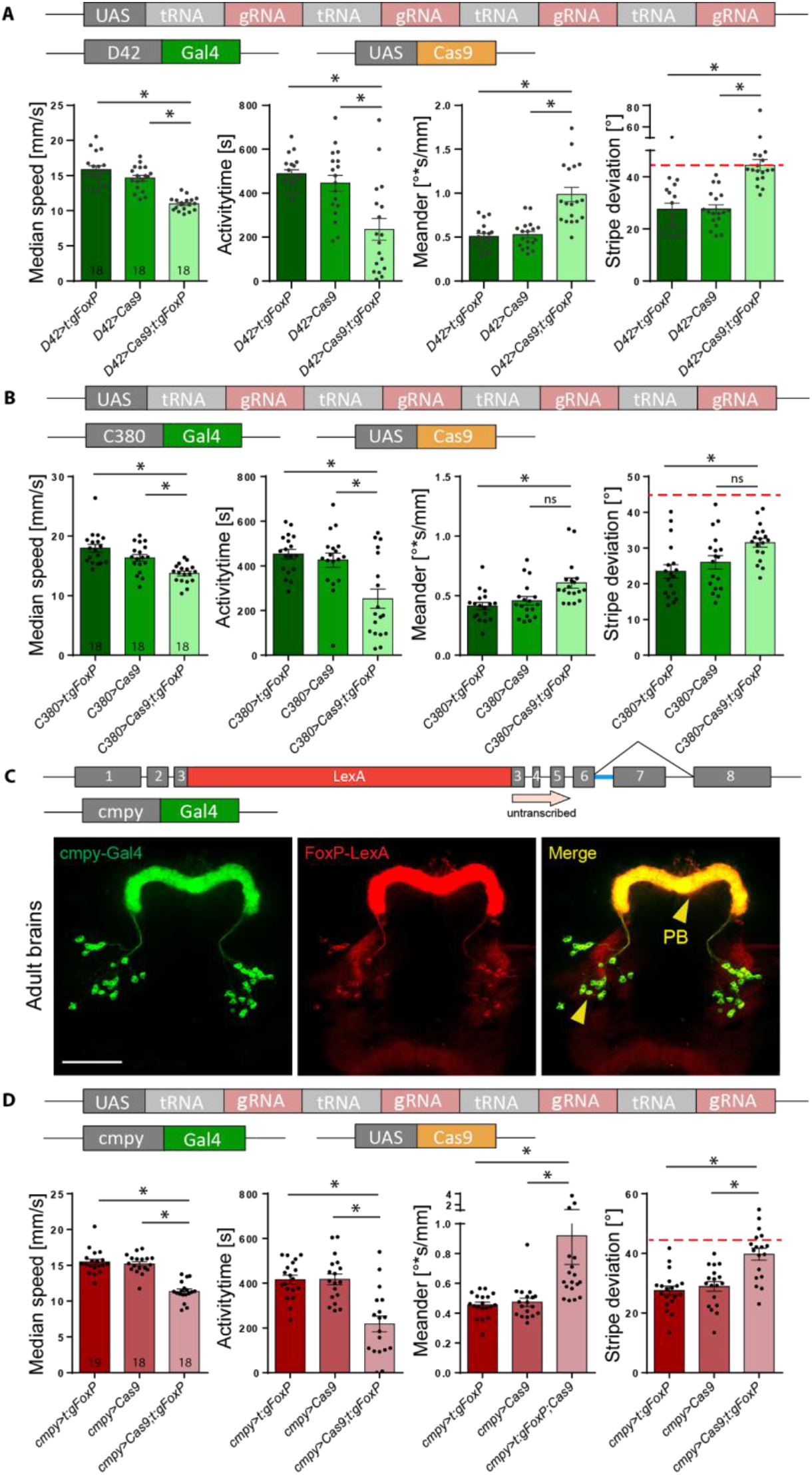
FoxP *is required in both motorneurons and protocerebral bridge for normal walking behavior in Buridan’s paradigm.* (A, B) Both motorneuron-specific driver lines (A) *D42-Gal4* (nonparametric tests for speed, activity time and meander) and (B) *C380-Gal4* show similar reductions in walking speed and activity time, but the spatial parameters fail to reach statistical significance despite trending in the same direction as in *D42*. (C) The *cmpy-Gal4* driver stains an overlapping set of protocerebral bridge (PB) neurons compared to our *FoxP-LexA* driver line. (D) Knocking out the *FoxP* gene in *cmpy-Gal4-positive* neurons leads to similar alterations in walking behavior in Buridan’s paradigm as a complete knock-out, i.e., reduced walking speed, reduced activity time, decreased stripe fixation and increased meander. Nonparametric test for meander. *p<0.005.

Perhaps the most prevalent *FoxP* expression can be observed in the protocerebral bridge (PB, Fig. 1). The driver line *cmpy-Gal4* targets the PB specifically and drives expression in *FoxP*-positive neurons (Fig. 10C). Removing the *FoxP* gene exclusively in these neurons led to a significant reduction of locomotor activity (Fig. 10D) as well as a reduction in stripe fixation and to more tortuous trajectories (Fig. 10E). In fact, the stripe deviation increased to an extent that it can no longer be distinguished statistically from a random walk at the 0.5% level (Wilcoxon signed rank test against 45°, V=33, p=0.02).

### 3.8. Conditional *FoxP-KO:* developmental stages

With *FoxP* being a transcription factor active throughout development and particularly important during pupal development [20,61], we knocked out *FoxP* in all neurons by adding the Gal4 repressor Gal80^ts^ to our panneuronal *FoxP-cKO* (Fig. 11A; raw data deposited at doi: 10.6084/m9.figshare.12607838) and treating the flies with a 48 hrs 30 °C heat treatment during the early pupal stage. This regime did not affect walking behavior in Buridan’s paradigm (Fig. 11B). Shifting the temperature treatment to immediately after eclosion also did not affect the flies’ behavior in Buridan’s paradigm (Fig. 11C). Taken together, these data indicate that *FoxP* is required for the proper development of, for instance, motor neurons and PB neurons, but once these circuitries are in place, FoxP expression does not appear to have any immediate mechanistic role in locomotion anymore.

**Fig. 11:**
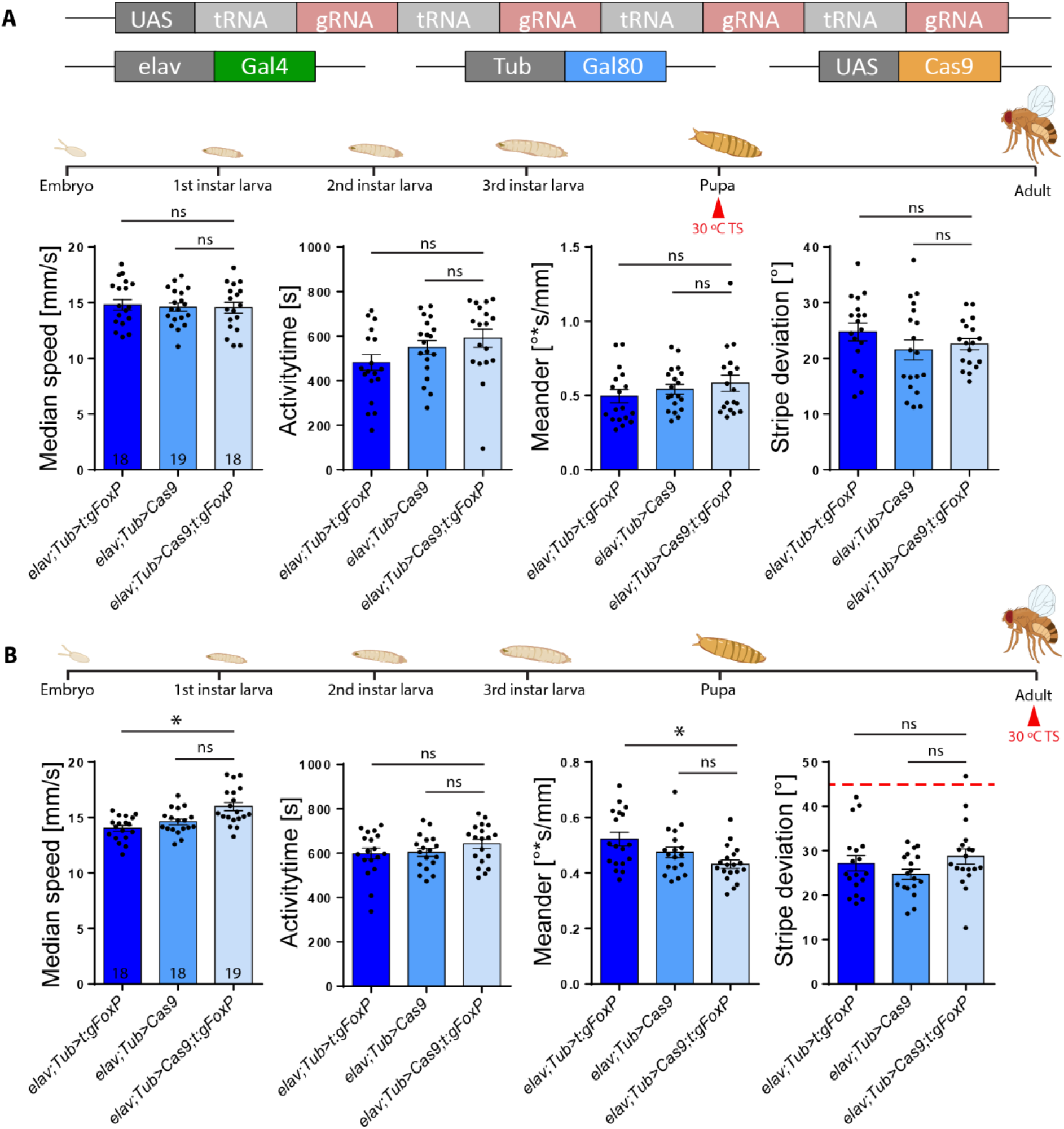
*Adult* FoxP *expression is not required for normal locomotor behavior in Buridan’s paradigm.* (A) Genetic tools used to perform the temporally controlled FoxP knock out. (B) Neither temporal (median speed and activity time) nor spatial (meander and stripe deviation) parameters are altered in adult flies in Buridan’s paradigm after inducing the FoxP KO in the early pupa. (C) Neither temporal nor spatial parameters are changed in adult flies in Buridan’s paradigm after inducing the FoxP KO immediately after eclosion. *p<0.005.

## 4. Discussion

We have edited the genomic locus of the *Drosophila FoxP* gene in order to better understand the expression patterns of the *FoxP* isoforms and their involvement in behavior. We have discovered that the isoforms differ with respect to their expression in neuronal tissue. For instance, we found isoform B (*FoxP-iB*) expression in neuropil areas such as the superior medial protocerebrum, the protocerebral bridge, the noduli, the vest, the saddle, the gnathal ganglia and the medulla, while areas such as the antennal lobes, the fan shaped body, the lobula and a glomerulus of the posterior ventrolateral protocerebrum contain other *FoxP* isoforms but not isoform B (summarized in Fig. 12) (raw data deposited at doi: 10.6084/m9.figshare.12607862). We also corroborated previous results [20] that *FoxP* is expressed in a large variety of neuronal cell types (Fig. 3). Our genomic manipulations created several new alleles of the *FoxP* gene which had a number of behavioral consequences that mimicked other, previously published alleles [20]. Specifically, we found that constitutive knockout of either *FoxP-IB* alone or of all *FoxP* isoforms affects several parameters of locomotor behavior, such as walking speed, the straightness of walking trajectories or landmark fixation (Figs. 6, 7). We discovered that mutating the *FoxP* gene only in particular neurons can have different effects. For instance, knocking *FoxP* out in neurons of the dorsal cluster (where *FoxP* is expressed) or in mushroom body Kenyon cells (where neither we nor[20] were able to detect FoxP expression) had no effect in Buridan’s paradigm (Fig. 9), despite these neurons being required for normal locomotion in Buridan’s paradigm [40,53,55]. In contrast, without *FoxP* in the protocerebral bridge or motor neurons, flies show similar locomotor impairments as flies with constitutive knockouts (Fig. 10). These impairments appear to be due to developmental action of the *FoxP* gene during larval development, as no such effects can be found if the gene is knocked out in all cells in the early pupal or adult stages (Fig. 11).

**Fig. 12:**
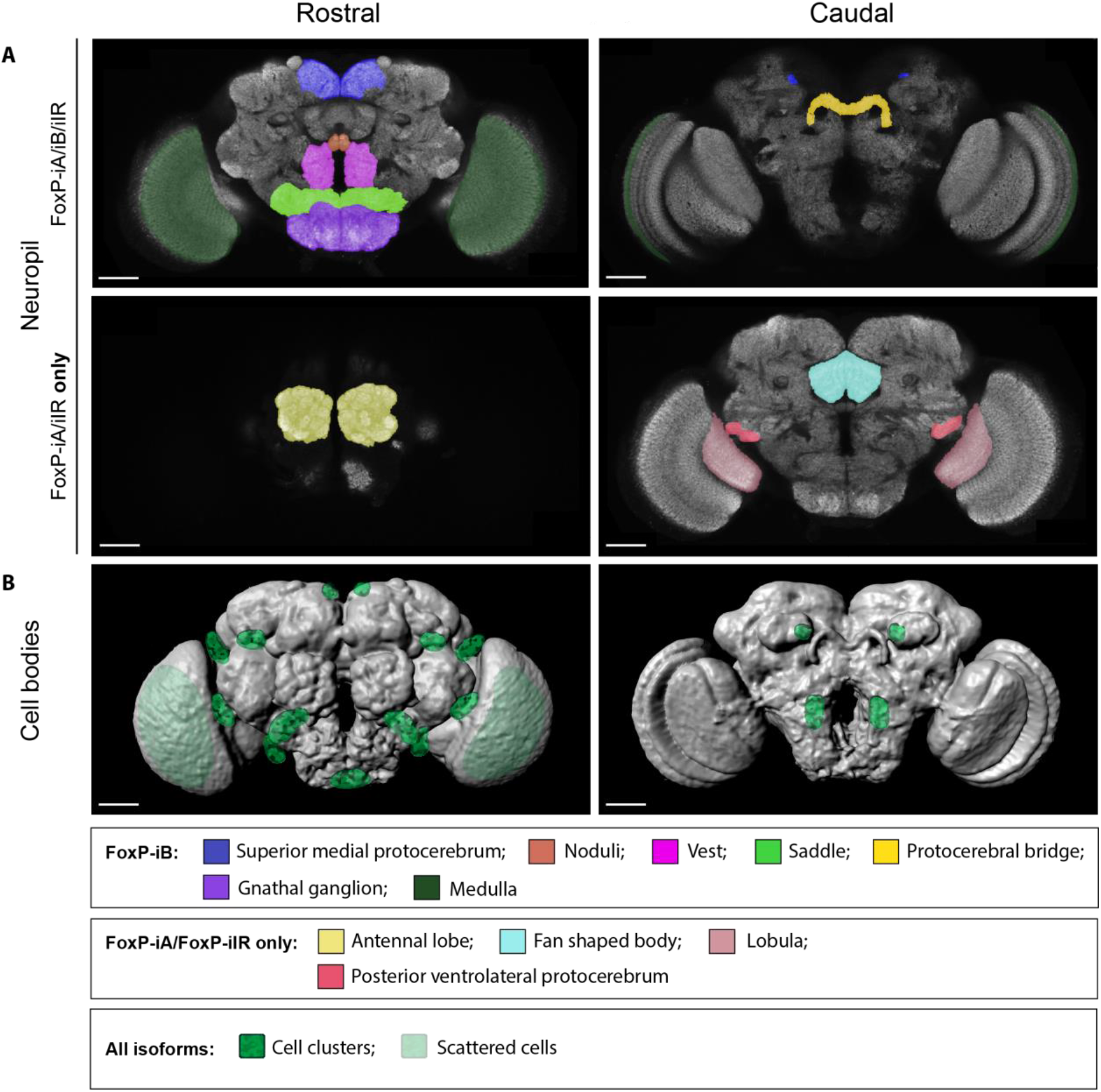
FoxP *expression pattern in the adult* Drosophila *brain*. (A) Rostral and caudal sections with neuropil areas marked for *FoxP-iB* expression (above) or for other *FoxP* isoforms excluding *FoxP-iB* (below). (B) Volume rendering of adult neuropil with marked approximate *FoxP*-positive cell body locations in the cortex. Scale bars: 50 μm.

### 4.1. *FoxP* expression

#### 4.1.1. Neuronal expression of *FoxP* is widespread but not in mushroom bodies

The exact expression pattern of *FoxP* has been under debate for quite some time now. Initial work combined traditional reporter gene expression with immunohistochemistry [21] (Table 4). Lawton and colleagues [21] created a *FoxP-Gal4* line where a 1.5 kb fragment of genomic DNA upstream of the *FoxP* coding region was used to drive Gal4 expression. These authors validated the resulting expression pattern with the staining of a commercial polyclonal antibody against FoxP. We used the same antibody in this work and observed perfect co-expression with our reporter (Fig. 1C). The Lawton *et al.* description of the *FoxP* expression pattern as a small number of neurons distributed in various areas of the brain, particularly in the protocerebral bridge (PB), matches our results and those of [20].

**Table 4:** The four previous reports and the present work describing FoxP expression patterns

| Publication | Method | Expression pattern |
| --- | --- | --- |
| Lawton et al. 2014 | - *Gal4* (1.5 kb fragment, pGaTB)<br>- antibody | - Small number of neurons in various areas of the brain<br>- not mushroom bodies |
| DasGupta et al. 2014 | *Gal4* (1.4 kb fragment, pBPGUw) | - Only mushroom bodies |
| Schatton and Scharff 2017 | *Gal4* (1.9 kb fragment, pBPGUw) | - Mushroom bodies and other areas |
| Castells-Nobau et al. 2019 | - GFP-tag (fosmid)<br>- antibody | - Abundant number of neurons in various areas of the brain<br>- not mushroom bodies |
| Present work | - genomic *Gal4*-tag (CRISPR/Cas9 HDR, pTGEM(0))<br>- antibody | - Abundant number of neurons in various areas of the brain<br>- not mushroom bodies |

Subsequent reports on FoxP expression patterns also used putative FoxP promoter fragments to direct the expression of Gal4 [34,61]. DasGupta and colleagues (2014) used a 1.4kb sequence upstream of the FoxP transcription start site, while Schatton and Scharff (2017) used 1.9 kb. Their larger fragment contained the sequences of the two previously used fragments (Table 4). In contrast to [20,21] and our work here, both studies reported expression in overlapping sets of mushroom body Kenyon cells. While DasGupta and colleagues implied that there was no expression outside of the mushroom bodies, Schatton and Scharff were not explicit. However, Schatton et al. independently reported that they had observed strong expression also outside of the mushroom bodies ([62] and pers. comm.).

The fourth and latest study reporting on *FoxP* expression in *Drosophila* [20] avoided the problematic promoter fragment method and instead tagged *FoxP* within a genomic segment contained in a fosmid [63], intended to ensure expression of GFP-tagged *FoxP* under the control of its own, endogenous regulatory elements. This study was the first to circumvent the potential for artifacts created either by selection of the wrong promoter fragment or by choosing an inappropriate basal promoter with the fragment (see below). However, since they also used insertion of a transgene, their expression pattern, analogous to that of a promoter fragment Gal4 line, may potentially be subject to local effects where the fosmid with the tagged *FoxP* was inserted. Castells-Nobau and colleagues (2019) also used the antibody generated by Lawton et al. (2014) to validate their transgenic expression patterns. The results reported with this technique are similar to those of Lawton et al. [21] and our work (summarized in Fig. 12B).

In an attempt to eliminate the last source of error for determining the expression pattern of FoxP in *Drosophila*, we used CRISPR/Cas9 with homology-directed repair to tag *FoxP in situ*, avoiding both the potential local insertion effects of the previous approaches and without disrupting the complex regulation that may occur from more distant parts in the genome. For instance, in human cells, there are at least 18 different genomic regions that are in physical contact with the *FOXP2* promoter, some of which act as enhancers [64]. The effects of these regions may be disrupted even if the entire genomic *FoxP* locus were inserted in a different genomic region as in [20]. Interestingly, the first promoter fragment approach [21] and the fosmid approach [20] agree both with our most artifact-avoiding genome editing approach and the immunohistochemistry with an antibody validated by at least three different *FoxP-KO* approaches. This converging evidence from four different methods used in three different laboratories suggests that *FoxP* is expressed in about 1800 neurons in the fly nervous system, of which about 500 are located in the ventral nerve cord. Expression in the brain is widespread with both localized clusters and individual neurons (Fig. 12B) across a variety of neuronal cell types. Notably, the four methods also agree that there is no detectable *FoxP* expression in the adult or larval mushroom bodies. In contrast, in honey bees, there is converging evidence of FoxP expression in the MBs [61].

#### 4.1.2. Understanding false-positive *FoxP* detection in mushroom bodies

This comparison of our data with the literature prompts the question why two different promoter fragment approaches [34,61] suggested *FoxP* expression in the mushroom bodies (confirmed by a ribosome-based approach, see below) when there is no FoxP protein detectable there.

A first observation is that Lawton *et al.* used the classic *hsp70*-based pGaTB vector [65] to create their Gal4 line, while both DasGupta *et al.* as well as Schatton and Scharff used the more modern *Drosophila* synthetic core promoter (DSCP)-based pBPGUw vector [66]. The two vectors differ with regard to their effects on gene expression. In addition to carrying two different basal promoters, the modern pBPGUw sports a 3’UTR that is designed to increase the longevity and stability of the mRNA over the pGaTB vector, which can result in two-fold higher Gal4 levels [67].

This observation is complemented by single-cell transcriptome data [68]. *FoxP* RNA can be detected in more than 4,100 brain cells, likely overcounting the actual FoxP expression more than 3-fold. For instance, *FoxP* RNA is detected in over 1,000 glial cells where none of the published studies has ever detected any FoxP expression (see also Fig. 2).

Taking these two observations together, it becomes plausible that there may be transient, low-level *FoxP* transcription in some mushroom body neurons (and likely thousands of other cells as well), which in wild type animals rarely leads to any physiologically relevant FoxP protein levels in these cells. Only when gene expression is enhanced by combining some arbitrary promoter fragments with genetically engineered constructs designed to maximize Gal4 yield such as the pBPGUw vector, such transient, low-abundance mRNAs may be amplified to a detectable level.

These considerations may also help explain why the ribosome-based method of [33] was able to detect *FoxP* RNA in mushroom body Kenyon cells: the transcript they detected may have been present and occupied by ribosomes, but ribosomal occupancy does not automatically entail translation [69]. It remains unexplained, however, how DasGupta *et al.* failed to detect all those much more strongly expressing and numerous neurons outside of the mushroom bodies. All of the above is consistent with other insect species showing FoxP expression on the protein level in their mushroom bodies [61], as only limited genetic alterations would be needed for such minor changes in gene expression.

The stochasticity of gene expression is a well-known fact [70–78] and known to arise from the transcription machinery [79]. Post-transcriptional gene regulation is similarly well-known [80–85]. It is thus not surprising if we observe that many cells often express transcripts that rarely, if ever, are translated into proteins. The final arbiter of gene expression must therefore remain the protein level, which is why we validated our expression analysis with the appropriate antibody. On this decisive level, FoxP has not been detected in the mushroom bodies at this point.

#### 4.1.3. Different *FoxP* isoforms are expressed in different neurons

Our genome editing approach allowed us to distinguish differences in the expression patterns of different FoxP isoforms. The isoform specifically involved in operant self-learning, *FoxP-iB,* is only expressed in about 65% of all *FoxP*-positive neurons. The remainder express either *FoxP-iA* or *FoxP-iIR* or both. Neurons expressing only non-iB isoforms are localized in the antennal lobes, the fan-shaped body, the lobula and a glomerulus of the posterior ventrolateral protocerebrum (Fig. 5). Combined with all three isoforms differing in their DNA-binding FH box, the different expression patterns for the different isoforms adds to the emerging picture that the different isoforms may serve very different functions.

### 4.2. *FoxP* and locomotor behavior

#### 4.2.1. *FoxP* is involved in both spatial and temporal parameters of walking

Alterations of *FoxP* family genes universally result in various motor deficits on a broad scale in humans [4,6,7,86] and mice [87–89] for both learned and innate behaviors. Also in flies, manipulations of the *FoxP* locus by mutation or RNA interference have revealed that *FoxP* is involved in flight performance and other, presumably inborn, locomotor behaviors [18,20–22] as well as in motor learning tasks [18].

The locomotor phenotypes described so far largely concerned the temporal aspects of locomotion, such as initiation, speed or duration of locomotor behaviors. Here, using Buridan’s paradigm [29,30], we report that manipulations of *FoxP* can also alter spatial aspects of locomotion, such as landmark fixation or the straightness of trajectories. Our results further exemplify the old insight that coarse assaults on gene function such as constitutive knock-outs of entire genes or isoforms very rarely yield useful, specific phenotypes [90]. Rather, it is often the most delicate of manipulations that reveal the involvement of a particular gene in a specific behavior. This fact is likely most often due to the pleiotropy of genes, often paired with differential dominance which renders coarse neurogenetic approaches useless in most instances, as so many different behaviors are affected that the specific contribution of a gene to a behavioral phenotype becomes impossible to dissect.

In the case of *FoxP,* it was already known, for instance, that the different isoforms affect flight performance to differing degrees [18] and that a variety of different *FoxP* manipulations affected general locomotor activity [20–22]. Here we show that a complete knock-out of either *FoxP-iB* or all isoforms affected both spatial and temporal parameters of locomotion, but the insertion mutation *FoxP^3955^* did not alter stripe fixation (Figs. 6, 7). Remarkably, despite the ubiquitous and substantial locomotor impairments after nearly any kind of *FoxP* manipulation be it genomic or via RNAi reported in the published literature, Das-Gupta et al. [34], failed to detect the locomotor defects of these flies.

While some of our manipulations did not affect locomotion significantly (e.g. knock-out in MBs or DCNs, Fig. 9, see below for discussion), most of them affected both spatial and temporal locomotion parameters (e.g., Figs. 6, 7, 9), despite these parameters commonly not co-varying [91]. Thus, while one would expect these behaviors to be biologically separable, our manipulations did not succeed in this separation.

#### 4.2.2. Locomotion does not require FoxP expression in all *FoxP* neurons

Taken together, the results available to-date reveal *FoxP* to be a highly pleiotropic gene with phenotypes that span both temporal and spatial domains of locomotion in several behavioral modalities, lifespan, motor learning, social behavior and habituation. It is straightforward to conclude that only precise, cell-type specific *FoxP* manipulations of specific isoforms will be capable of elucidating the function this gene serves in each phenotype. With RNAi generally yielding varying levels of knock-down and, specifically, with currently available *FoxP* RNAi lines showing only little, if any, detectable knock-down with RT-qPCR (data deposited at doi: 10.6084/m9.figshare.12607667 and Annette Schenck, pers. comm.), CRISPR/Cas9-mediated genome editing lends itself as the method of choice for this task. Practical considerations when designing multi-target gRNAs for *FoxP* prompted us to begin testing the CRISPR/Cas9 system as an alternative to RNAi with an isoform-unspecific approach first, keeping the isoform-specific approach for a time when we have collected more experience in this technique. In a first proof-of-principle, we used CRISPR/Cas9 to remove *FoxP* from mushroombody Kenyon cells (MBs), dorsal cluster neurons (DCNs), motor neurons (MNs) and the protocerebral bridge (PB).

MBs have been shown to affect both spatial and temporal aspects of locomotion [e.g., 50,51-59] and Castells-Nobau et al. [20] reported a subtle structural phenotype in a subset of MB Kenyon cells that did not express *FoxP.* As detailed above, two groups have reported *FoxP* expression in the MBs and it appears that some transcript can be found in MB Kenyon cells. With a substantial walking defect both in *FoxP^3955^* mutant flies (which primarily affects *FoxP-iB* expression [18]) and in flies without any *FoxP* (Fig. 7), together with the MBs being critical for normal walking behavior, the MBs were a straightforward candidate for a cell-type-specific *FoxP-KO*. However, flies without *FoxP* in the MBs walk perfectly normally (Fig. 9C). There are two possible reasons for this lack of an effect of our manipulation: either FoxP protein is not present in MBs or it is not important in MBs for walking. While at this point we are not able to decide between these two options, our expression data concurring with those from previous studies [20,21] suggest the former explanation may be the more likely one (see also above). Remarkably, a publication that did report *FoxP* expression in the MBs [34] did not detect the walking deficits in *FoxP^3955^* mutant flies despite testing for such effects. Motor aberrations as those described here and in other *FoxP* manipulations [20–22,35] constitute a potential alternative to the decision-making impairments ascribed to these flies in DasGupta *et al.* [34].

DCNs were recently shown to be involved in the spatial component (landmark fixation) of walking in Buridan’s paradigm [40], but removing *FoxP* from DCNs showed no effect (Fig. 9B), despite abundant *FoxP* expression in DCNs (Fig. 9A). It is possible that a potential effect in stripe fixation may have been masked by already somewhat low fixation in both control strains. On the other hand, even at such control fixation levels, significant increases in stripe deviation can be obtained (e.g., Fig. 10A). Before this is resolved, one explanation is that *FoxP* is not required in these neurons for landmark fixation in Buridan’s paradigm, while the neurons themselves are required.

MNs are involved in all aspects of behavior and have been shown to be important for operant self-learning [60]. With abundant expression of *FoxP* in MNs (Fig. 3A), we considered these neurons a prime candidate for a clear *FoxP-cKO* phenotype. Indeed, removing FoxP specifically from MNs only, mimicked the effects of removing the gene constitutively from all cells (Fig. 10A, B). It is noteworthy that this manipulation alone was sufficient to affect both temporal and spatial parameters, albeit only one of the two driver lines showed clearcut results (D42). One would not necessarily expect motorneurons to affect purportedly ‘higher-order’ functions such as landmark fixation. It is possible that the higher tortuosity in the trajectories of the flies where D42 was used to drive our UAS-gRNA construct is largely responsible for the greater angular deviation from the landmarks in these flies and that this tortuosity, in turn is caused by the missing FoxP in motorneurons. Alternatively, D42 is also driving in non-motorneurons where *FoxP* is responsible for landmark fixation. The driver line C380 showed similar trends, albeit not quite statistically significant at our alpha value of 0.5%, suggesting that potentially the increased meander parameter may be caused by motorneurons lacking *FoxP*.

The PB is not only the arguably most conspicuous *FoxP*-positive neuropil, it has also been reported to be involved in temporal aspects of walking [92–94]. Moreover, the PB provides input to other components of the central complex involved in angular orientation [22,95–97]. Similar to the results in MNs, removing *FoxP* from a small group of brain neurons innervating the PB, phenocopies constitutive *FoxP* mutants.

Taken together, the MN and PB results suggest that both sets of neurons serve their locomotor function in sequence. At this point, it is unclear which set of neurons precedes the other in this sequence.

#### 4.2.3. *FoxP* is only required during larval development to ensure normal locomotion

There is ample evidence that the *FoxP* family of transcription factors acts during development in a variety of tissues [9,98,99]. What is less well known is if adult *FoxP* expression serves any specific function. A recent study in transgenic mice in operant conditioning and motor learning tasks showed postnatal knock-out of FOXP2 in cerebellar and striatal neurons affected leverpressing and cerebellar knock-out also affected motor-learning [89]. At least for these tasks in mammals, a *FoxP* family member does serve a postnatal function that is independent of brain development (brain morphology was unaltered in these experiments). Also in birds, evidence has been accumulating that adult *FoxP* expression serves a song plasticity function [100–105]. Our temporally controlled experiments (Figs. 8, 11) suggest that at least locomotion in Buridan’s paradigm can function normally in the absence of *FoxP* expression in the adult, as long as *FoxP* expression remains unaltered during larval development. Future research on the role of *FoxP* in locomotion and landmark fixation hence needs to focus on the larval development before pupation.

## Acknowledgements

Technical assistance was provided by Klara Krmpotić (Buridan experiments with *FoxP-iB-Gal4*) and Marcela Loza. We are indebted to David Deitcher for sharing his FoxP antibody with us. We are grateful to three anonymous reviewers for their insightful suggestions, which significantly increased the quality of the manuscript. This project was funded by the DFG (Deutsche Forschungsgemeinschaft, grant BR 1892/17-1).

## Data sharing

All original raw data are deposited with figshare (http://figshare.com), references in the text.

## Author contributions

Conceived and designed the experiments: OP, MR, BB. Performed the experiments: OP. Analyzed the data: OP, MR, BB. Contributed reagents/materials/analysis tools: MR, BB. Wrote the paper: OP, BB. Initiated the project: BB. Critically revised the manuscript: OP, MR, BB. The authors declare no existing conflicts of interest.

